# Resistance to targeted therapies as a multifactorial, gradual adaptation to inhibitor specific selective pressures

**DOI:** 10.1101/504837

**Authors:** Robert Vander Velde, Nara Yoon, Viktoriya Marusyk, Arda Durmaz, Andrew Dhawan, Daria Myroshnychenko, Diego Lozano-Peral, Bina Desai, Olena Balynska, Jan Poleszhuk, Liu Kenian, Mohamed Abazeed, Omar Mian, Aik Choon Tan, Eric Haura, Jacob Scott, Andriy Marusyk

## Abstract

Despite high initial efficacy, therapies that target oncogenic kinases eventually fail in advanced, metastatic cancers. This failure in initially responsive tumors is the direct result of the evolution of drug resistance under therapy-imposed selective pressures. In contrast to the massive body of experimental research on the molecular mechanisms of resistance, understanding of its evolutionary origins and dynamics remains fragmented. Using a combination of experimental studies and mathematical modeling, we sought to dissect the evolution of resistance to different clinical ALK inhibitors in an experimental model of ALK positive NSCLC. We found that resistance can originate from heterogeneous, weakly resistant, sub-populations with variable sensitivity to different ALK inhibitors. Instead of the commonly assumed stochastic single hit (epi) mutational transition, or drug-induced reprogramming, we found evidence of a hybrid scenario, of gradual, multifactorial development through acquisition of multiple cooperating genetic and epigenetic adaptive changes, amplified by selection. Additionally, we found that intermediate resistance phenotypes might present unique, temporally restricted collateral sensitivities, absent in therapy naïve or fully resistant cells, suggesting new opportunities for therapeutic interference.

## INTRODUCTION

Despite inducing strong clinical responses, inhibitors that target abnormal activities of oncogenic tyrosine kinases (TKIs), including front and second line TKIs directed against EML4-ALK fusion oncogene in non-small cell lung cancers (NSCLC), are rarely curative in advanced disease^1^. Since ALK targeting TKIs (ALK-TKIs) typically fail to eradicate all of the tumor cells, the survivors can evolve resistance-conferring phenotypic adaptations, culminating in the outgrowth of resistant subpopulations and eventual clinical relapse. Though we know that tumors change and evolve under drug-imposed selective pressures, this knowledge is not accounted for in the standard of care in clinical practice, where tumors are treated as static entities, with therapy switched only after relapse. Adequate understanding of the evolutionary trajectories and underlying dynamics might offer an opportunity to drastically improve clinical outcomes with therapeutic tools that are already at our disposal^2–5^. For example, by accounting for frequency dependent selection and fitness tradeoffs a pilot clinical trial in castration resistant prostate cancer significantly extended progression free survival while reducing side effects of the treatment, using different scheduling of standard therapy drugs^6^.

In contrast to massive research efforts that have advanced our understanding of the molecular mechanisms of resistance, evolutionary dynamics during the acquisition of resistance remains understudied. Consequently, our conceptual understanding of how resistance evolves, which inform mathematical modeling studies, is often based on conjectures from mechanistic studies, rather than direct experimental inquiry suitable for informing quantitative models. For example, demonstrations of the pre-existence of therapy-associated molecular alterations in sub-populations of tumor cells within treatment naïve samples have led to the widely prevalent assumption that resistance to targeted therapies arises simply due to a competitive release of pre-existent genetically (or epigenetically) distinct, therapy-resistant subpopulations under drug-imposed selective pressures^7, 8^. On the other hand, a growing body of experimental studies suggest that therapy resistance can emerge *de novo* from drug-tolerant persister (DTP) cells^9^, capable of surviving in the drugs and limited proliferation, but incapable of supporting robust tumor growth. DTPs are often assumed to constitute a phenotypically well-defined sub-population, and sometimes equated with cancer stem cells^10^. While DTP cells cannot sustain positive net tumor growth in the face of therapy, they can maintain residual disease and serve as a substrate for mutational or epigenetic conversions to therapy-resistant phenotypes, capable of driving robust growth rates in the face of therapy^9^. Finally, as therapies can induce adaptive phenotypic changes on shorter time scales, independent of selection for more fit phenotypes, development of resistance has also been viewed from a differentiation/reprograming paradigm^11–13^.

Motivated by our interest in developing evolutionary informed therapeutic scheduling to tackle TKI resistance, we decided to investigate the origin and evolutionary dynamics that underlie the rapid development of therapy resistance of an *in vitro* model of ALK+ NSCLC, the patient derived H3122 cell line. We found that, upon exposure to different ALK-TKIs, NCI-H3122 cells rapidly and predictably develop strong drug resistance. Resistance originates *de novo*, from weakly resistant heterogeneous sub-populations, which differ in fitness when exposed to different ALK-TKIs. Levels of resistance gradually increase under therapy, through acquisition of multiple cooperating genetic and epigenetic mechanisms, through TKI-specific phenotypic trajectories. In contrast to therapy naïve or fully resistant cells, these evolving populations show strong collateral sensitivity to the dual EGFR/HER2 inhibitor lapatinib, suggesting a temporally restricted opportunity to interfere with the development of resistance.

## RESULTS

### I. Continuous exposure to ALK-TKIs leads to acquired resistance mediated by predictably distinct phenotypes

To understand the development of acquired resistance to ALK-TKIs, we took advantage of a well-characterized ALK+ NSCLC cell line, NCI-H3122, which can rapidly develop resistance to multiple clinically relevant ALK-TKIs using dose escalation protocols^14^. Using resistant cell lines from our previous study^14^, we continued dose escalation, eventually selecting for cells capable of growing in high, clinically relevant concentrations of the drugs (up to 1 µM crizotinib, 4 µM lorlatinib, 2 µM alectinib, 200 nM ceritinib). H3122 cells with **e**volved **r**esistance to ALK-TKI (erALK-TKI) displayed strong collateral resistance toward all other tested ALK-TKIs, with 5-100x higher IC50 (**Fig. 1A** and **Table S1**). Despite the similar shift in IC50 in all of the tested ALK-TKIs, erALK-TKI cells developed through selection by different inhibitors displayed more divergent sensitivities to higher drug concentrations. Consistent with the clinical efficacy of alectinib and lorlatinib as a second line therapy after failure of crizotinib^15, 16^, high concentrations of alectinib and lorlatinib strongly inhibited cells with evolved resistance to crizotinib (erCriz) and ceritinib (erCer). The resistance was at least partially heritable, as the erALK-TKI phenotypes were partially or completely maintained after drug holiday (**Fig. S1A**). Further, upon xenograft transplantation into immunocompromised NSG mice, cells with **e**volved **r**esistance to alectinib (erAlec) and lorlatinib (erLor), formed tumors capable of maintaining net growth when challenged with clinically relevant concentrations of alectinib and lorlatinib, despite the 3 week gap between tumor implantation and initiation of the treatment (**Fig. 1B, C**).

**Fig. 1.**
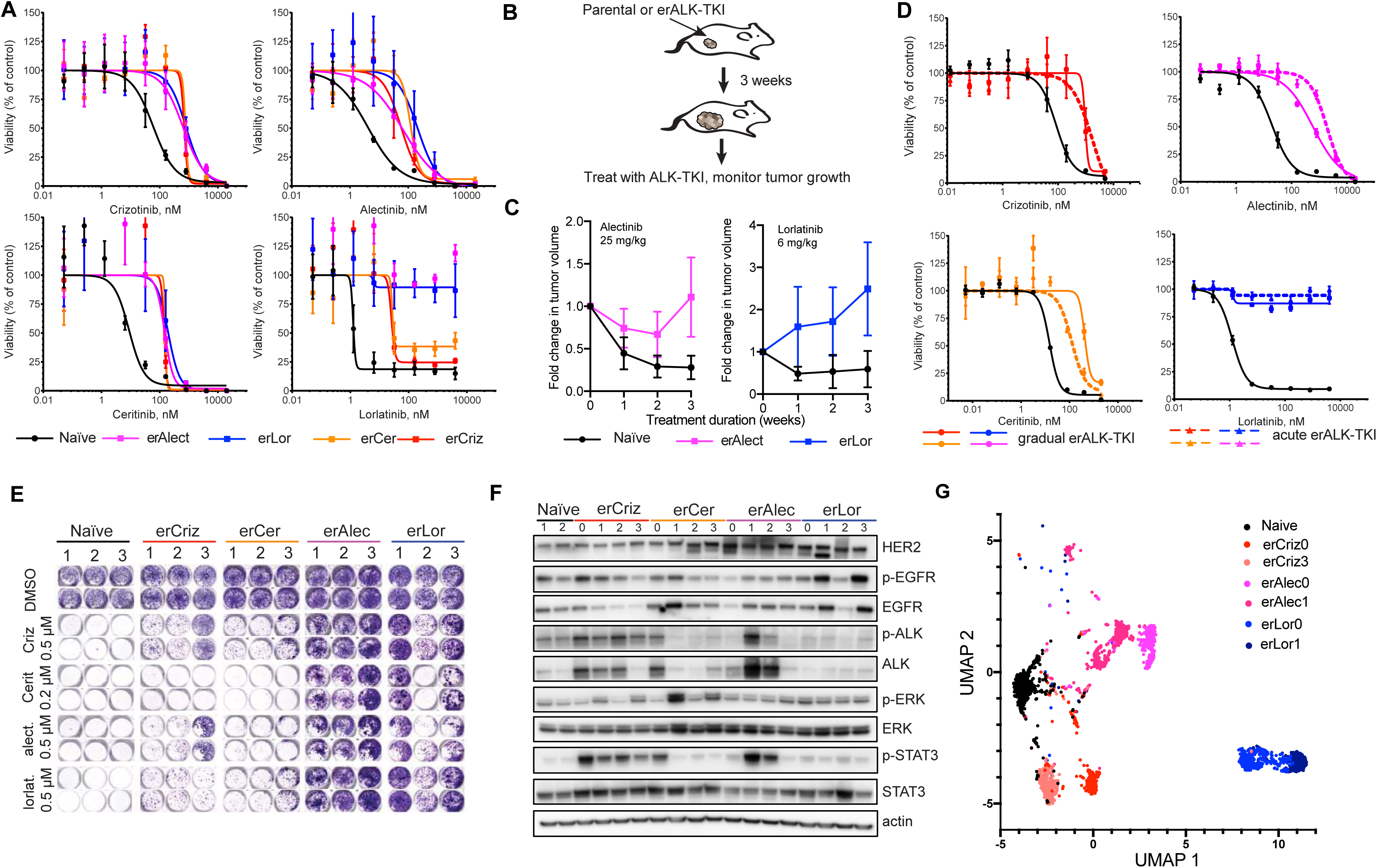
Characterization of evolved resistance to distinct ALK-TKIs. **A.** Sensitivities of the treatment naïve H3122 cell line and the indicated erALK-TKI derivative cell lines to the indicated ALK inhibitors, measured by Cell Titer Glo assay. Error bars show standard deviation. **B**. Experimental schemata of the xenograft experiment. **C**. Growth dynamics of xenograft tumors produced by inoculation of the indicated cells. Treatment with indicated doses of ALK TKIs or vehicle control is started 3 weeks post implantation. Error bars show standard deviation. **D**. Comparison of resistance levels of erALK-TKI cells evolved by gradual and acute drug exposure, measured by Cell Titer Glo assay. Error bars show standard deviation. **E**. Sensitivity of independent derivates of erALK-TKI cells to the indicated ALK-TKIs, measured by crystal violet assay. **F**. Immunoblot analysis of the expression levels of the indicated proteins in the independent derivations of erALK-TKI cells. “0” denotes gradually derived erALK TKI cell lines, presented in Figure A, “1-3” indicates independent derivate sub-lines obtained by acute selection for resistance in high drug concentrations, same as in E. **G.** UMAP analysis of single cell RNA-seq expression of the indicated cell lines.

Whereas drug dose escalation protocols are commonly used in studies to identify resistance mechanisms^17^, they do not reflect the development of resistance in clinics, as patients are treated upfront with the highest tolerated dose of the drugs. Therefore, we asked whether resistance can develop in the face of acute exposure to clinically relevant concentrations of ALK-TKIs. To this end, we plated treatment naïve H3122 cells in the presence of 0.5 µM crizotinib, 2 µM alectinib and lorlatinib or 100 nM ceritinib, replating cultures as needed to relieve contact growth inhibition of the surviving cells. After 3-4 months of this selection, we obtained rapidly proliferating erALK-TKI cell lines, with resistance levels comparable to those achieved with the dose escalation protocol (**Fig. 1D**).

We next asked to what extent the differences in cross-sensitivities of erALK-TKI cells towards different ALK inhibitors are attributable to the choice of specific ALK inhibitor. To answer this, we independently derived triplicate lines for each of the ALK-TKIs used in this study, with the acute exposure protocol, and compared their cross-sensitivities to higher concentrations of the inhibitors. Consistent with previous findings (**Fig. 1A**), erLor and erAlec cells demonstrated stronger resistance to high concentrations of different ALK-TKIs, compared to erCriz and erCer cell lines (**Fig. 1E**). Similar to the resistant cell lines derived by gradual exposure, resistant phenotypes in cell lines derived by acute exposure to ALK-TKIs were largely heritable (**Fig. S1B**).

Given the surprising predictability in ALK-TKI cross-sensitivity phenotypes, we interrogated the extent of predictability of changes in EML4-ALK dependent signaling pathways. Immunoblot analyses, performed on cells cultured in the absence of the drugs for 48 hrs (to reduce the direct impact of ALK-TKIs on cell signaling) revealed that all of the erCriz cell lines displayed increased phosphorylation of EML4-ALK, in most cases in association with elevated EML4-ALK protein levels, as well as increased STAT3 phosphorylation (**Fig. 1F**). In contrast, all of the erLor cell lines lacked the increase in STAT3 phosphorylation and displayed reduced ALK phosphorylation. Instead, they expressed higher levels of EGFR, HER2, or both. erAlec and erCer cell lines exhibited more diverse (less predictable) phenotypes. To assess the phenotypes more globally, we examined mRNA expression levels of 230 cancer-related genes using Nanostring nCounter GX human cancer reference panel. Principal component and hierarchical clustering analyses for mRNA expression were largely consistent with the immunoblot evaluation of phosphorylation (**Fig. S2A, B**). erCer, evolved through acute selection, as well as acutely and gradually evolved erCriz and erLor cells formed distinct clusters; in contrast, phenotypes of erAlec were more diverse.

Potentially, observed differences could reflect either cell population-wide phenotypic changes or differences in relative abundance of distinct pre-existent subpopulations. To discriminate between these two possibilities, we performed single cell RNA sequencing of erALK-TKI cell lines, focusing on two cell lines per specific ALK-TKI, with the highest divergence in PCA analysis of NanoString data (**Fig. S2A**). Uniform manifold approximation and projection (UMAP)^18^ dimension reduction of single cell expression data revealed relative phenotypic homogeneity within individual ALK-TKI lines, with a high degree of similarity among cell lines derived with the same inhibitor (**Fig. 1G**). Thus, despite a degree of stochasticity, acquired resistance to specific ALK-TKIs is associated with highly convergent, inhibitor-specific phenotypes.

### II. Resistance to distinct ALK TKIs develops from pre-existing heterogeneous weakly resistant sub-populations

Evolution of acquired resistance to targeted therapy is often assumed to reflect a simple expansion of therapy resistant sub-populations^19^. Whereas the observation that predictable distinctions between resistance associated phenotypes are selected in different ALK-TKIs contradicts this notion, we decided to interrogate the pre-existence of fully resistant phenotypes more directly. To this end, we seeded treatment naïve NCI-H3122 cells at clonogenic densities in the presence of ALK-TKIs or DMSO (vehicle control). At 10 days post seeding, the majority of treatment naïve cells plated in DMSO, as well as erALK-TKI cells plated in DMSO or ALK-TKIs, formed macroscopic colonies. In contrast, whereas treatment naïve cells formed multiple microscopic colonies, we could not detect colonies with comparable sizes to the large colonies formed by resistant cells, upon plating as many as 10,000 treatment naïve cells in the presence of ALK-TKIs (**Fig. 2A**). At the same time, limiting dilution experiments revealed that 1:338 - 1:660 of treatment naïve NCI-H3122 cells give rise to robustly growing drug-resistant colonies within 10 weeks of drug exposure (**Fig. 2B, C**). Whereas these observations cannot exclude pre-existence of rare (<0.01%) fully resistant sub-populations, acquired resistance, within our experimental system, likely originates from weakly resistant intermediates, capable of survival and limited proliferation under the drugs. This inference is consistent with time-lapse microscopy examination of the dynamics of drug resistance emergence, where robust growth in the presence of ALK-TKIs was observed, but only after significant delay (**Suppl. video 1**).

**Fig. 2.**
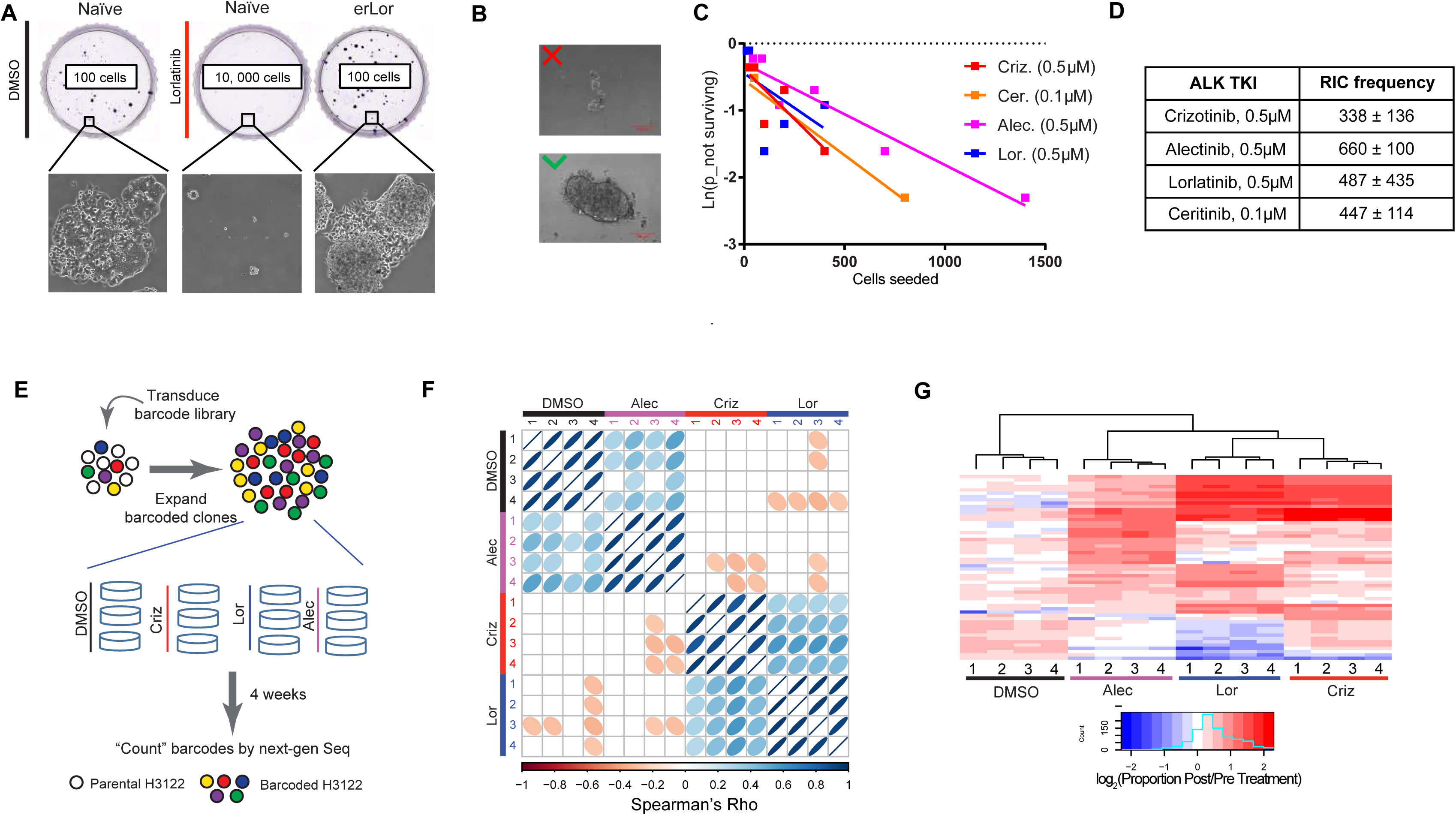
Resistance to ALK TKIs arises from heterogeneous DT sub-populations. **A.** Representative images of crystal violet stained whole plates and microscopic images of colonies (at 10x magnification) following 10 days of culture. **B**. Illustration of the size cut-off criteria used for quantitation of limiting dilution experiments. **C**. **D**, Fitting of limiting dilution assay data and quantitation of the frequency of resistance-initiating cells. **E**. Schemata of the clone tracing experiment. **F**. Spearman pairwise correlation analysis of barcodes, which have reached frequency, exceeding the highest frequency barcode in the initial cell mix (0.0077), from the indicated biological replicates. **G**. Hierarchical clustering analysis of the same “winning” barcodes as in **F**.

The existence of a distinct weakly resistant phenotypic state which is a precursor to *bona fide* resistance, termed “tolerance” or “persistence” has been widely studied in microbiology ^20, 21^. Based on the observation of a similar phenomenon in the context of response to TKIs, the term “drug tolerant persisters” (DTPs) has been introduced to describe a weakly resistant subpopulation in EGFR+ NSCLC^9^, and later in other cancers^13, 22, 23^. DTPs are commonly viewed as a main contributor to minimal residual disease^24^. Potentially, DTP cells can reflect a distinct population that either pre-exists prior to therapy, or arise, either stochastically or deterministically, in response to drug-induced stress. To discriminate between these scenarios, we used tracing with selectively neutral DNA barcodes, an approach which has been previously used to demonstrate pre-existence of sub-populations resistant to EGFR inhibitors in EGFR+ lung cancers^25^. We transduced H3122 with a high complexity lentiviral ClonTracer library at a low multiplicity of infection (MOI) to ensure that most of the transduced cells are labelled with a single unique barcode. Following the elimination of non-transduced cells with puromycin selection, and ∼100x expansion of the barcoded cells, we took a baseline aliquot, then separated the cells into parallel quadruplicate cultures, and exposed them to 0.5 µM alectinib, lorlatinib, and crizotinib or DMSO control. After 4 weeks of incubation, barcode frequencies were determined by next generation sequencing, and compared to the baseline frequencies (**Fig. 2E**). Clear evidence of both negative and positive selection was observed in all treatment groups (including DMSO controls), as barcode diversity, captured with Shannon diversity index, decreased (**Fig. S3A**), while several subpopulations have expanded (**Fig. S3B**). Spearman ranking of barcodes, which increased in frequency above the highest barcode frequency observed in the baseline sample, indicated a strikingly high degree of correlation between biological replicates within the same treatment condition, indicating pre-existence of relatively stable weakly resistant subpopulations (**Fig. 2F**). However, correlation between samples treated with different ALK TKIs was either absent or much less pronounced, indicating that selective pressures exerted by different ALK TKIs might amplify distinct pre-existing weakly resistant sub-populations. Unsupervised hierarchical clustering analysis revealed a partial overlap between “winning” clones (barcodes) across multiple ALK TKIs, indicating that at least some of the distinctions reflect quantitative rather than qualitative differences, and some of the pre-existent phenotypes were equally adaptable to different ALK TKIs (**Fig. 2G**). Thus, resistance to an ALK-TKI in H3122 cells originates from pre-existent, heterogeneous, subpopulations, with weak resistance and varying sensitivity to distinct ALK-TKIs.

In contrast to the fitness cost of classical persistence under baseline growth conditions, considered to be a form of bet hedging^21^, clones enriched in ALK-TKIs, on average, were also slightly enriched when cultured in the absence of the drugs (DMSO control) (**Fig. S3C**).

### III. Gradual development of ALK-TKI resistance

Having established that resistance originates from heterogeneous pre-existing tolerant populations, we next asked how it progresses toward full resistance. The prevalent assumption in the modeling, experimental and clinical communities is that resistance results from a single-hit, transition mediated by acquisition of a genetic mutation or by a (meta) stable change in expression of resistance-conferring genes through a stochastic or drug-induced epigenetic switch^7, 26^. To test whether resistance in H3122 cells develops through a single hit transition, we employed the following clonogenic assay (**Fig. 3A**). Therapy naïve cells were pre-cultured in the presence of ALK-TKIs (crizotinib or lorlatinib) for 1-3 weeks, then harvested and seeded at clonogenic densities in the presence or absence of the drug. After 7 days of culture, we measured clonogenic survival as well as the sizes of the resulting colonies. Since prolonged exposure to the drugs is expected to eliminate sensitive populations, while enriching for drug tolerant and resistant ones, we expected the clonogenic proportion to increase over time. Indeed, the clonogenic proportion in 0.5 µM crizotinib progressively increased from the initial 2.6% to 26% at week three, while the clonogenic proportion in 0.5 µM lorlatinib increased from the initial 1% to 17% at week three (**Fig. 3B**). Consistent with the lack of proliferation penalty in the barcoding experiments, we did not observe substantial changes in size of colonies, formed by cells with evolved or evolving resistance, in the absence of the drugs (**Fig S4A, B**). If the transition from tolerance to resistance were to result from a single-hit mutational or epigenetic switch, we would expect to observe a bimodal distribution of colony sizes, representing tolerant and resistant colonies, with the fraction of the latter gradually increasing (**Fig. 3A**). As expected, longer ALK TKI exposure lead to an increase in the average colony size. However, instead of the expected bimodal size distribution, this increase was apparently homogeneous, suggesting a gradual development of resistance (**Fig. 3C**). Analysis of the colony size distributions at the intermediate (2 and 3 weeks) time-points using Kolmogorov-Smirnov statistics revealed that the observed colony sizes cannot be explained by mixing colony size distributions of tolerant and erALK-TKI cells (**Fig. S4C, D**).

**Fig. 3.**
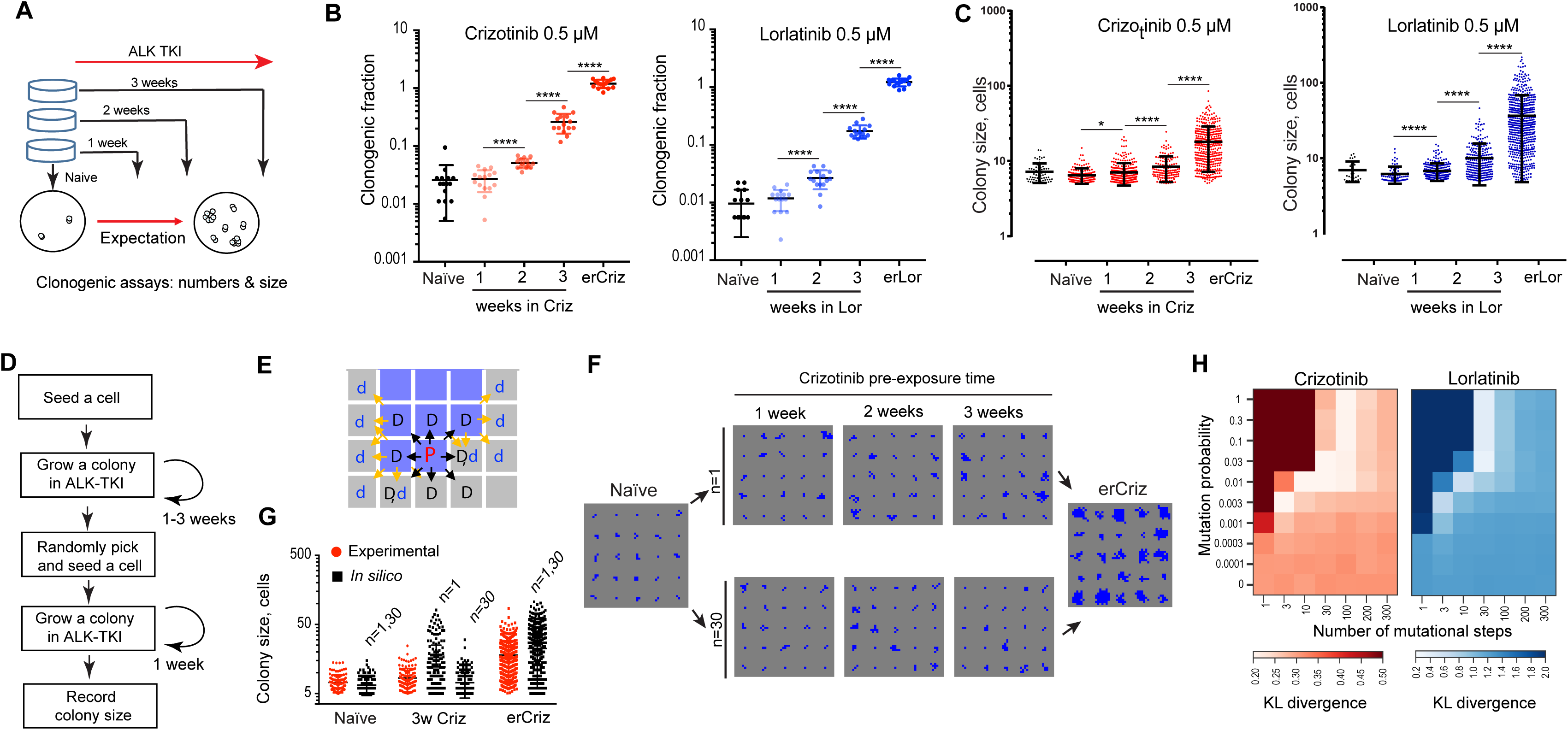
Gradual development of acquired ALK TKI resistance. **A.** Schemata of the experiment to assess the mode of progression from tolerance to resistance. **B.** Clonogenic survival in the presence of the drug after pre-incubation for indicated times in crizotinib and lorlatinib; data is normalized to clonogenic survival in DMSO control. Error bars show standard deviation. **C.** Distributions of colony sizes in the presence of the indicated ALK TKI. * and **** represent p-values of a Mann-Whitney non-parametric test, <0.05 and <0.0001 respectively. Error bars show standard deviation. **D.** Logical flow diagram for the agent-based mathematical model. The model simulates growth from initial seeding through pre-incubation and the clonogenic assay. **E.** Proliferation space check scheme. Blue and grey cells denote occupied and empty spaces respectively. “P” stands for the parent cell, “D” for daughter cell, “d” for displaced cell. Black arrows indicate options for placement of daughter cells, yellow arrows indicate options for displaced cells. Proliferation can occur if a nearby space is either immediately available, or separated by a single cell, in which case this cell is pushed into an empty space, with an extra copy of the proliferating cell displacing it. **F.** Example of colony growth simulations, initiated from cells pre-incubated in crizotinib for the indicated time, during the final colony assay, contrasting n=1 versus n=30. Error bars show standard deviation. **G**. Comparing divergence between *in silico* and experimental data, with n=1 and 30. Error bars show standard deviation. **H.** Kullback–Leibler divergence-based comparison of the experimental data with the outcomes of simulations, covering parameter space for the indicated mutation probabilities and numbers of mutational steps.

Inferences with Kolmogorov-Smirnov statistics might be obfuscated by the fact that (epi)mutations mediating transition from tolerance to *bona fide* resistance can potentially occur not only during the 1-3 weeks growth preceding the clonogenic assay, but also during the clonogenic stage of the assay. Therefore, to discriminate between a single step and multi-step development of resistance, we developed an agent-based mathematical model of the evolution of therapy resistance, which enabled *in silico* simulation of the experimental assay across a range of mutational probabilities and numbers of mutational steps (**Fig. 3D**, **Mathematical Supplement**). This model simulates growth of individually seeded cells and their progeny in the presence/absence of the drug both prior and during the clonogenic assay. Cells are seeded into a 2d lattice (simulating the surface of a culture dish). If space is available (no more than one cell separating the cell from an empty space), a cell can proliferate with a given probability inferred from the experimental data (**Fig. 3E**). To account for the variability in observed colony sizes, the maximal proliferation probability is set for each simulated colony individually, based on random sampling of sizes of colonies produced by resistant cells. For the initial proliferation probability, we use one single value calibrated with the whole distribution of naïve cell colony sizes, assuming minor variability in the proliferation rates of such cells. At each cell division, a cell can increase in division rate, reflecting adaptive (epi-)mutations. Cells can transition from the initial to maximal division probability by either a single step (n=1), or multiple (n>1) steps, representing fractional increments of the single transition (**Fig. 3D** **and Mathematical Supplement**).

Whereas the initial and maximal growth rates are fixed based on the experimental data, the intermediate rates within the *in silico* simulations are not. They can differ based on two parameters: a) (epi) mutational probability and b) the number of mutational steps required to reach maximum division probability (n). Since we do not know the (epi) mutation probability, we explored the outcomes with the full range of possible values (0-1), and the number of mutational steps ranging from 1 to 300. With each choice of parameters, we generated 100,000 random *in silico* simulations (**Fig. 3F** and **Suppl. video 2)**. Differences between *in vitro* and *in silico* colony size distributions were assessed using Kullback-Leibler divergence. We found that a single mutational step provided the poorest fit to the data for all mutation probabilities. The best fit was achieved with a number of mutational steps in the range of 3-100 (**Fig. 3G, H**). Notably, inclusion of death rates, bi-directional change in cell fitness and consideration of mutations that combine multiple increments at a single cell division did not qualitatively change this outcome (**Fig. S4 E, F** **and Mathematical Supplement**). Therefore, the experimentally observed dynamics of the development of ALK TKI resistance is inconsistent with a transition toward resistance occurring exclusively through a single step transition. Instead, our analyses suggest a gradual evolution through incremental increases in drug resistance. Of note, since the improvements could be achieved through multiple increments within a single cell division, our analyses do not exclude a possibility of a hybrid scenario, where, in rare cases, resistance can be acquired in a single step.

**Fig. 4.**
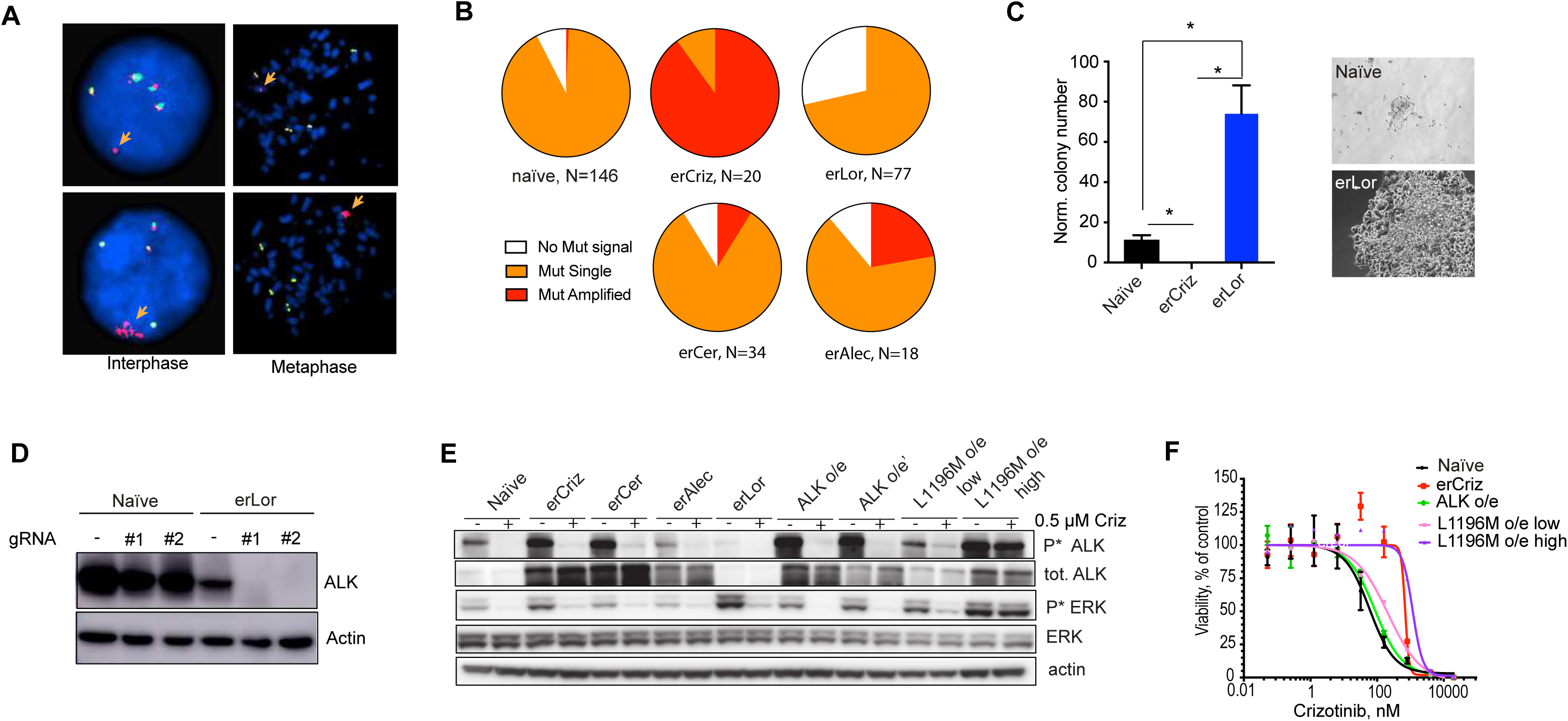
Contribution of ALK expression changes to erALK-TKI phenotypes. **A.** Representative images for interphase and metaphase FISH analysis for EML4-ALK fusions and amplification status. Separation of 3’ (red) probe from 5’ (green) probe indicates ALK fusion event (orange arrows). **B**. Frequency of cells with the indicated EML4-ALK fusion and amplification status in the gradually evolved erALK-TKI cell lines. **C**. Impact of CRISPR mediated genetic ablation of ALK on clonogenic survival of the indicated H3122 derivates. * indicates p< 0.05 in a two-tail unpaired t-test. **D**. Evaluation of EML-ALK ablation status by immunoblotting analysis. **E**. Immunoblot evaluation of the expression and activity of EML4-ALK oncogenic signaling in the indicated cell lines with evolved and engineered resistance. **F**. Impact of retrovirally mediated overexpression of “wild type” EML4-ALK and its L1196M mutant variant on sensitivity to crizotinib, measured by Cell Titer Glo assay. Error bars show standard deviation.

These inferences of graduality of acquisition of ALK TKI resistance are inconsistent with a prevalent idea of acquired resistance attributable to a single genetic or epigenetic mechanism. Since molecular mechanisms of resistance to ALK-TKIs have been extensively characterized, including numerous studies in H3122 experimental system, we decided to interrogate the involvement of previously identified ALK-TKI resistance mechanisms in erALK-TKI phenotypes, and their ability to mediate single-hit acquisition of resistance.

Clinical resistance to ALK inhibitors is frequently associated with point mutations in the kinase domain of ALK, which reduce drug binding^27^. However, hotspot mutations were absent in the erALK-TKI cell lines, as revealed by targeted Sanger sequencing of PCR-amplified cDNA (**Fig. S5**). Given that clinical ALK-TKI resistance is often associated with EML4-ALK amplification, and that we observed an increase in the expression of EML4-ALK in some of the erALK-TKI resistant cell lines (**Fig. 1F**), we tested the amplification status of EML4-ALK in treatment naïve, as well as in representative erALK TKI resistant cell lines (lines “0” in **Fig. 1F**) using the mutational break apart FISH assay. The majority of treatment naïve H3122 cells displayed 4 copies of the wild type allele and one copy of the fusion allele, with a minor sub-population where the fusion gene signal could not be detected. Some of the erALK-TKI cells displayed multiple copies of the fusion allele, consistent with gene amplification (**Fig. 4A**). Extrachromosomal amplification of oncogene-containing DNA has been recently implicated in the rapid evolution of TKI resistance^28^, however examination of metaphase spreads revealed that the amplified alleles were localized within the chromosome. Consistent with the immunoblotting results shown in **Fig. 1F**, we observed substantial heterogeneity in the amplification status of EML4-ALK, both between and within erALK-TKI cell lines (**Fig. 4B**). The majority of erCriz cells, as well as a fraction of erAlec and erCer cells displayed EML4-ALK amplification. In contrast, erLor cells not only lacked EML4-ALK amplification, but also the proportion of oncogene negative cells was significantly higher (p<0.0001 in a Chi-square test) than in the treatment naïve cells, suggesting that *loss of EML4-ALK* might have been selectively advantageous under the more potent ALK-TKI.

We next investigated whether differences in EML4-ALK copy numbers corresponded to differences in oncogene dependence. We transfected treatment naïve, erCriz and erLor cells with constructs expressing Cas9 and two different guide RNAs targeting ALK and selected for puromycin resistant colonies. No colonies could be observed for erCriz cells, suggesting a critical dependency on EML4-ALK (**Fig. 4C**). Parental cells formed few small colonies, resembling tolerant colonies formed upon exposure to an ALK-TKI (**Fig. 1D**). In contrast, erLor cells formed multiple colonies that displayed no evidence of growth inhibition (**Fig. 4C**), despite complete ablation of the protein expression of EML4-ALK gene (**Fig. 4D**). Interestingly, puromycin resistant naïve cells transfected with guide RNA directed against ALK displayed lack of EML4-ALK ablation, consistent with strong selection for variants that uncouple antibiotic resistance from guide RNA expression, indicating a strong selective disadvantage of losingEML4-ALK expression. This observation is consistent with reduced baseline EML4-ALK expression in erLor cells (**Fig. 1F**) and suggests a conversion into a phenotype that is independent of EML4-ALK oncogenic activity.

Given that EML4-ALK amplification, resulting in overexpression is considered to provide a *bona fide* therapy resistance mechanism^29^, we asked whether the observed increase in EML4-ALK expression could explain the resistance observed in the erALK-TKI cell lines with EML4-ALK amplification. Retroviral overexpression of EML4-ALK resulted in protein levels of total and phosphorylated EML4-ALK which closely resemble those observed in EML4-ALK amplified erALK-TKI cells (**Fig. 4E**). Further, after exposure to crizotinib, cells with retrovirally overexpressed EML4-ALK retained residual levels of ALK phosphorylation similar to those observed in the erALK-TKI cells (**Fig. S6A**). Despite this, EML4-ALK overexpressing cells displayed only a marginal increase in crizotinib resistance (**Fig. 4F**), suggesting that while ALK amplification contributes to resistance, it is insufficient to fully account for it.

Given the insufficiency of EML4-ALK overexpression to fully account for resistance, we decided to interrogate the functional impact of the most common resistance-associated point mutation, L1196M, which is thought to provide a single hit resistance mechanism to crizotinib^29^. Surprisingly, at low expression levels, achieved with <1 retroviral MOI, L1196M expression only moderately decreased crizotinib sensitivity (**Fig. 4G**). In contrast, higher overexpression of the mutant protein, at levels similar to those observed with cells containing amplified “wild type” fusion gene, achieved with high MOI, blocked the ability of crizotinib to shut down phosphorylation of EML4-ALK, as well as its main downstream effector ERK (**Fig. 4E, S6A, B**), and provided a resistance level that is similar or higher to that observed in erCriz cells (**Fig. 4F**). Whereas the results with L1196M overexpression are consistent with previously reported sufficiency to confer resistance in NIH-H3122 cells^30, 31^, and ALK mutations do co-occur with EML4-ALK amplification^32^, we are not aware of reports of amplification of ALK mutant alleles. Thus, common resistance-associated mutational changes might be insufficient to provide a single hit solution to the challenge posed by ALK-TKI induced selective pressures.

Next, we examined resistant cell lines for the presence of additional copy number alterations. Whereas Oncomine Focus Assay^33^ failed to detect additional common mutations, CytoScanHD SNP array revealed additional genetic changes, including recurrent chromosomal amplifications in chromosomes 2, 3, 12 and 17. Interestingly, 2 out of 3 examined erLor lines contained Chr12 p12.1-p11.1 amplification, containing KRAS, whereas a third one contained Chr1 p13.2-p12 amplification containing NRAS (**Fig. S7A**). Notably, chromosomal amplification of genomic regions containing KRAS and NRAS were associated with elevated expression of these proto-oncogenes (**Fig. S7B**).

Given the growing evidence for the importance of recurrent (semi)heritable non-genetic changes in gene expression in resistance to targeted and cytotoxic therapies, we used RNA-Seq analysis to examine changes in gene expression after a 48 hr drug holiday (to reduce the direct impact of ALK TKI on gene expression). We found multiple gene expression changes, previously implicated in TKI and chemotherapy resistance, including increased expression of multiple RTKs (EGFR, HER2, FGFR, AXL, EPHA2, etc.), cytokines, ECM & ECM receptors and other types of molecules, suggesting a complex, multifactorial nature of resistance (**Fig. 5A**). Importantly, co-expression analysis of cell lines in the CCLE database revealed that the genes with upregulated expression in erALK-TKI cells belonged to distinct gene co-expression clusters, suggesting that the resistance-associated changes in gene expression are unlikely to represent a single coordinated transcriptional program switch (**Fig. S8A, B**). Notably, gene set enrichment analysis revealed several shared enriched gene sets (**Fig. S8C**). In particular, all of the examined resistant cell lines displayed an epithelial to mesenchymal transition (EMT) signature. Whereas EMT has been described as ALK-TKI resistance mechanism^34, 35^, our results suggest a need for a more nuanced interpretation, given the differences in specific resistance associated molecular changes observed in different erALK-TKI cell lines, which would be missed under the EMT umbrella term.

**Fig. 5.**
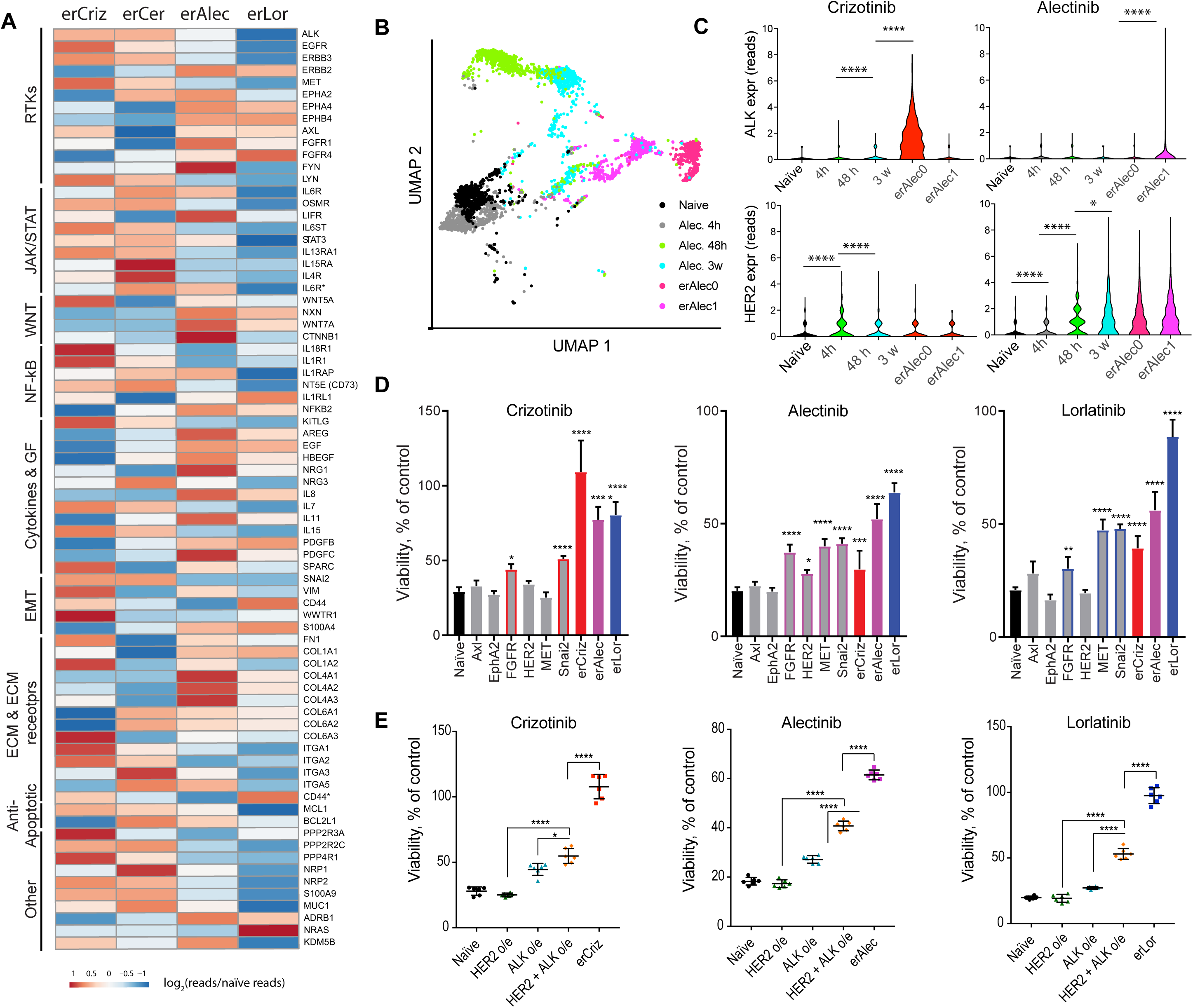
Evolved resistance to ALK TKI integrates multiple mechanisms. **A.** Elevated mRNA expression for the indicated genes, previously associated with chemotherapy or targeted therapy resistance, in erALK-TKI cell lines across the indicated functional categories. **B**. UMAP analysis of single cell mRNA expression data from cells exposed to 0.5 μM alectinib for the indicated time duration. **C**. Violin plot of expression levels of ALK and HER2 following indicated duration of exposure to the indicated ALK-TKIs. *, **, ***, **** represent p<0.05, p<0.01, p<0.001 and p<0.0001 of a Mann-Whitney U-test. **D.** ALK-TKI sensitivity of engineered cell lines, lentivirally overexpressing individual putative resistance associated expression changes to the indicated ALK-TKIs (0.5 μM), determined by Cell Titer Glo assay. Error bars show standard deviation. **E.** Impact of combination of individual resistance mechanisms toward sensitivity to indicated ALK-TKIs (0.5 μM), determined by Cell Titer Glo assay. *, **, ***, **** represent p<0.05, p<0.01, p<0.001 and p<0.0001 of ANOVA analysis with Dunnett’s (B) or Tukey’s (C) multiple comparison correction. Error bars show standard deviation.

To gain further insights into the dynamics of phenotypic changes during the evolution of resistance, we analyzed phenotypic changes following different exposure times to ALK TKIs using single cell expression profiling. Notably, UMAP analysis revealed gradual phenotypic progression from naïve to resistant phenotypes (**Fig. 5B, S9**). Even brief (4 hours) alectinib exposure substantially impacted cell phenotypes, suggesting that acquisition of resistance might reflect not only the action of drug-imposed selection, but also direct drug-induced cell adaptation. On the other hand, some of the naïve cells mapped to a major phenotypic cluster of one of the erCriz (erCriz3) samples (**Fig. S9**), consistent with the notion that some of the resistant phenotypes might arise from selection of pre-existent sub-populations. To gain further insight into the temporal dynamics of acquisition of resistance-conferring expression changes, we analyzed the expression of ALK and HER2 at single cell levels, at different time points, after exposure to an ALK-TKI. Consistent with high levels of ALK expression and genetic amplification of EML4-ALK in erCriz cell lines, expression of ALK increased upon exposure to crizotinib, but not alectinib. In contrast, expression of HER2 became elevated within 48 hours of exposure to both drugs, suggesting a direct adaptive response, then decreased upon prolonged incubation with crizotinib, while staying up-regulated or increasing further under alectinib (**Fig. 5C**).

To explore a possible epigenetic mechanism underlying the observed early and heritable changes in gene expression associated with ALK-TKI exposure, we analyzed the global repatterning of Histone 3.3 lysine 27 acetylation (H3K27ac), a post-translational histone modification associated with strong enhancer elements. We performed ChIP-Seq analysis to characterize differential H3K27ac distribution at gene regulatory elements in naïve and evolved resistant lines. We found that different ALK-TKI resistant phenotypes are associated with distinct global patterns of H3K27 acetylation (**Fig. S10A**). Changes in gene expression in the resistant cell lines were associated with changes in H3K27 acetylation (**Fig. S10B**). Consistently, we found new H3K27ac peaks in the vicinity of genes upregulated in individual ALK-TKI treated lines, e.g. ERBB2 / HER2 (**Fig. S10C**). Consistent with the differences in protein expression (**Fig. 1F**) the new peaks were observed in erAlec and erLor lines, but not in the erCriz lines. Interestingly, new peaks were also observed for genes with chromosomal genomic amplification in relevant cell lines (EML4, K-RAS and N-RAS) (**Fig. 12D**), suggesting the involvement of both genetic and epigenetic mechanisms in the elevated expression of putative drivers of resistance. Taken together, these data suggest that stable gene expression changes, in resistance-associated genes, might be established and maintained by a combination of genetic and epigenetic changes.

To evaluate the functional significance of elevated expression of the resistance-associated genes on ALK-TKI sensitivity, we tested the impact of overexpression of selected genes, previously implicated in TKI resistance. We found that lentiviral overexpression of HER2, FGFR, AXL, and SLUG significantly increased resistance to multiple ALK-TKIs (**Fig. 5D**). However, similar to the insufficiency of EML-4 ALK amplification to fully account for ALK-TKI resistance, resistance levels observed in these engineered cell lines fell short of levels observed in the erALK-TKI lines, suggesting that evolved resistance might reflect the combined effect of multiple contributing changes. To directly evaluate this possibility, we interrogated the impact of the combination of individual genetic and transcriptional changes. We combined overexpression of EML4-ALK, which is amplified in many resistant cell lines (**Fig. 1F**) and provides a very modest decrease in crizotinib sensitivity (**Fig. 4F**), with expression of HER2 (at lower levels compared to overexpressing cells used for **Fig. 5B**), which is also elevated in multiple erALK-TKI cell lines (**Fig. 1F**) and can provide modest resistance to different ALK inhibitors, when overexpressed at high levels (**Fig. 5D**). Whereas HER2 overexpression had very limited impact on ALK-TKI sensitivity, it significantly increased resistance conferred by EML4-ALK overexpression, though still failing to recapitulate resistance levels observed in erALK-TKI cells (**Fig. 5E**). These results support the notion that the evolution of resistance proceeds through acquisition of multiple genetic and epigenetic changes, that, under inhibitor-specific selective pressures of ALK-TKIs, shape resistant phenotypes in a combinatorial fashion.

### IV. Evolving resistance is associated with temporally restricted fitness tradeoffs

It is frequently assumed that, according to the principle of an evolutionary fitness tradeoff^36^, resistance-conferring phenotypes should be associated with strong fitness penalties compared to drug naïve tumor cells, when outside of the primary treatment^6, 37–39^. This fitness penalty could enable evolutionary informed adaptive therapy, based on creating a “tug of war” between therapy sensitive and therapy resistant populations by sequencing primary drug with strategic treatment breaks, informed by tumor’s response to the drugs^4^. Thus, we examined growth of erALK-TKI cell lines in the presence and absence of the drugs. Whereas in some cases, cells taken off the drugs indeed proliferated slower than drug naïve control, some of the resistant cell lines proliferated at similar or even higher rates (**Fig. S11A**). Interestingly, all of the examined erCriz and 1 out of 3 of the examined erAlec cell lines displayed higher rates of proliferation in the presence of the inhibitors, consistent with previously reported observations in melanoma^40, 41^. The remaining cell lines were either modestly inhibited, or unaffected by the ALK-TKI used for their selection (**Fig. S11A**).

Inferences obtained from *in vitro* 2d cultures, however, could be misleading. The fitness of tumor cells is context dependent^42, 43^, and *in vitro* 2d cultures do not capture death/proliferation dynamics within tumors *in vivo*. Further, non-cell autonomous interactions between phenotypically distinct subpopulations could significantly alter fitness in competitive settings^44, 45^. Therefore, we tested the fitness impact of resistance conferring adaptations by implanting mixtures of differentially labelled (GFP and mCherry) resistant and naïve cells into subcutaneous tumors or into the orthotopic environment of the lungs (via tail vein injections). Only erCriz cells injected into the lungs displayed signs of reduced fitness compared to therapy naïve cells (**Fig. S11B**). Interestingly, erLor cells had a significant selective advantage over naïve cells both in subcutaneous and orthotopic tumor contexts. These data suggest that lower fitness outside of the TKI is not an obligatory fitness tradeoff, and that acquisition of resistance might be associated with increased fitness in the absence of treatment.

Growth out of the primary therapy is just one of many potential contexts where an evolutionary tradeoff might be manifested. Altered context, created by application of a different drug, to which phenotypes with acquired resistance to ALK-TKI are collaterally sensitive, could provide an alternative strategy to expose cells to a resistance-linked fitness tradeoff^46, 47^. Consistent with the up-regulation of HER2 in alectinib resistant cells, H3122 cells failed to develop resistance to alectinib in the presence of dual EGFR/HER2 inhibitor lapatinib (**Fig. S12A**). However, erAlec cells were not sensitized to lapatinib as a single agent, and only some of the independently derived erAlec cell lines were inhibited by the combination of alectinib and lapatinib (**Fig. S12B**). To examine this apparent discrepancy, we examined the sensitivity of intermediate stages of alectinib resistance to lapatinib. We found that evolving cells were remarkably sensitive to lapatinib, even as a single agent, although this sensitivity gradually diminished as cells became more resistant to alectinib (**Fig. 6A, B**). Notably, this sensitivity cannot be explained by pre-existence of lapatinib sensitive, alectinib tolerant sub-populations, as treatment with 10 µM lapatinib for up to 3 weeks did not substantially impact sensitivity of H3122 to alectinib (**Fig. S12C**). Consistently, administration of lapatinib as a single agent, following an initial three weeks in alectinib, or as a combination therapy, was able to achieve complete elimination of H3122 cells *in vitro* (**Fig 6C**).

**Fig. 6.**
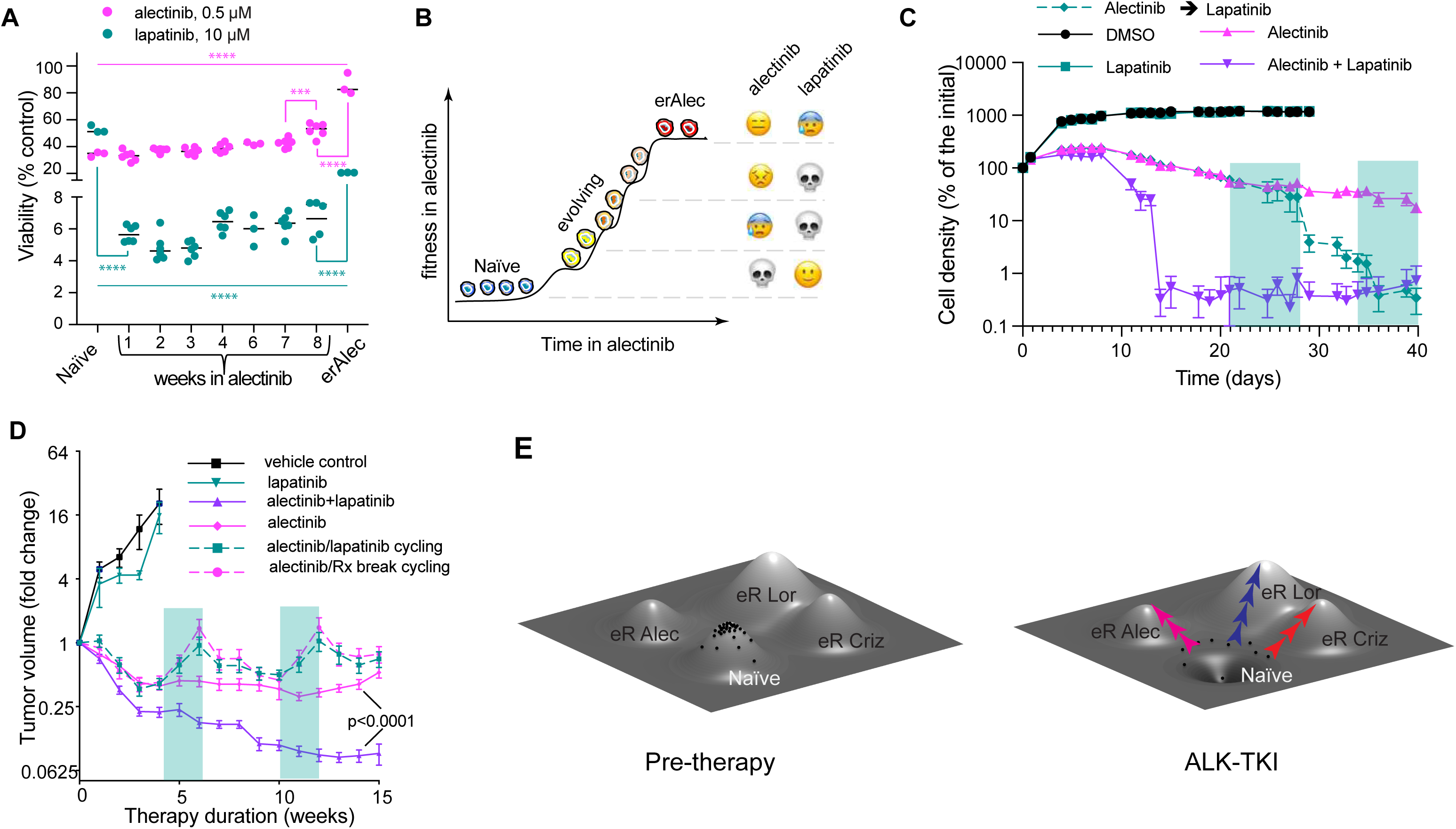
Gradual evolution of resistance provides a new therapeutic window for collateral sensitivity. **A.** Differential lapatinib sensitivity of H3122 cells following indicated exposure time to 0.5 µM alectinib toward alectinib or lapatinib, measured by Cell Titer Glo assay. One-way ANOVA was used for both drugs (p<0.0001). Adjacent timepoints were compared using Sidak’s multiple comparison tests (*** and **** represent p<0.001 and p<0.0001). Error bars show standard deviation. **B**. Schemata: evolving resistance to alectinib is associated with collateral sensitivity to lapatinib. **C**. Lapatinib can prevent the development of resistance to alectinib *in vitro* both as a combination treatment, or in drug cycling. Green shading indicates switching from alectinib to lapatinib monotherapy. Residual signal in the combination therapy and cycling groups reflects autofluorescence as visual examination revealed lack of surviving tumor cells. **D**. Change in volume of H3122 xenograft tumors treated with indicated therapies; treatment initiated 3 weeks post tumor implantation. Green shading indicates switching from alectinib to lapatinib monotherapy or vehicle control. **** represents p<0.001 of the interaction term of a repeated measurements 2-way ANOVA. Error bars indicate SEM. **D**. Evolving resistance interpreted with a fitness landscape metaphor. Naïve cells occupy a local fitness peak. Drug exposure reshapes the landscape, turning this fitness peak into a fitness trough. Different ALK inhibitors act on partially distinct outliers, directing their evolution toward distinct fitness peaks.

Encouraged by this observation, we asked whether alectinib/lapatinib cycling or a combination treatment with the two drugs could outperform alectinib monotherapy *in vivo*. We found that lapatinib was completely ineffective in adaptive cycling with alectinib, as tumor relapse during lapatinib cycles was indistinguishable from the relapse observed with a drug holiday (**Fig. 6D**). Most likely, this lack of efficiency *in vivo* reflects its inability to reach the high concentrations required to achieve collateral sensitivity *in vitro*. Still, lapatinib significantly increased tumor sensitivity to alectinib in a combination therapy setting (**Fig. 6D**). Whereas, at this point, the clinical utility of our observations remains unclear, they provide a proof of principle that temporally restricted collateral vulnerabilities of evolutionary intermediates might be exploited therapeutically to improve responses, and potentially, to forestall the emergence of resistance.

## Discussion

Despite substantial advances in deciphering the molecular mechanisms of resistance to TKI based targeted therapies, and the development of more effective drugs, advanced “targetable” lung cancers remain incurable, as tumors eventually acquire resistance and relapse. Developing strategies to interfere with evolving resistance has the potential to substantially improve long term survival outcomes. However, this is contingent on a correct understanding of the underlying evolutionary dynamics. In contrast to the large body of experiment-derived knowledge on individual molecular mechanisms of resistance, our understanding of its evolutionary causes and dynamics remains less developed. The subject of evolutionary-informed therapies has received significant attention from mathematical modelers^48–51^. However, given the paucity of experimental data acquired to derive solid assumptions for building models, these modeling studies often have to rely on conjectures from reductionist, mechanism-centered studies, thus limiting their potential.

Acquired resistance is commonly viewed through two major conceptual frameworks. According to the first, currently dominant gene-centric framework, resistance arises from a selective expansion of genetically or epigenetically distinct (meta)stable sub-populations. The subpopulation(s) could either pre-exist therapy or arise *de novo* from mutational conversion of sensitive or tolerant cells. In either case, clinical and experimental studies operating within this paradigm typically reduce acquired resistance to a single cause, such as a point mutation (*e.g.* the L1196M gatekeeper mutation^52^), gene amplification (e.g., EML4-ALK^52^, cMET^53^, etc.), or a non-mutational stable change in expression of a resistance-conferring gene (e.g., IGFR^54^). Consequently, the assumption of a single hit resistance mechanism is highly prevalent in the mathematical modelling community^26, 49, 51^.

The second framework views resistance as the result of drug-induced reprogramming, where phenotypic plasticity enables tumor cells to directionally “rewire” their signaling, metabolic and gene-expression networks to cope with inhibitor-induced perturbations^13, 55, 56^. In this system-biology based framework, resistance can be gradual and multifactorial, however the causal role of selection is sometimes rejected due to its link with the mutation-centric paradigm^13^.

The two frameworks are not necessarily viewed as strictly mutually exclusive, as they can be bridged in a two-step process involving reprogramming mediated formation of DTP sub-populations followed by an (epi) mutational switch to full resistance. Our results, however, suggest an alternative scenario of gradual, multifactorial resistance, which integrates features of the two paradigms described above. Based on i) the observation of gradual development of resistance from heterogeneous weakly resistant sub-populations, ii) the co-occurrence of both genetic and non-genetic changes, of which at least some appear to be directly induced by drug exposure, iii) the insufficiency of single resistance-associated mechanisms to confer full resistance, and iv) the additive action of individual resistance-associated mechanisms, we propose the following model: Selective pressures imposed by therapies act on phenotypic heterogeneity, stemming from both stochastic (genetic and epigenetic) and drug-induced changes, leading to a gradual increase in population fitness through acquisition of additional genetic and epigenetic changes until a local fitness peak is reached (**Fig. 6F**). We speculate that different ALK inhibitors lead to different evolutionary trajectories, due to differences in both selective pressures and direct drug-induced phenotypic changes (owing to different on and off target inhibitory activities). Of note, this proposed framework essentially mirrors Darwin’s original concept of evolution by natural selection, as a gradual adaptation to environment-imposed selective pressures, rather than the currently prevalent mutation-centric re-interpretation of Darwin’s idea within the cancer research community^57^.

At this point, generalizability of this proposed framework beyond the H3122 experimental model remains to be explored. However, gradual, multifactorial acquisition of resistance has been recently observed in an elegant experimental study of acquired BRAF resistance in melanoma^13^. The authors rejected the Darwinian explanation based on the lack of pre-existent resistance and thus interpreted their observations strictly within a “reprogramming” paradigm. We argue that their data is highly consistent with our model, which does not reduce Darwinian selection to genetically distinct pre-existing variants. Likewise, a recent single cell sequencing study in triple negative breast cancer demonstrated that while resistance to chemotherapy can be traced to preexistent genetically distinct subpopulations, resistant cells have acquired new phenotypic changes, most likely reflecting epigenetic modlifications^58^.

The proposed concept of gradual resistance does not exclude the possibility of pre-existence of full resistance, or a mixed scenario, where tumor relapse might reflect a combined contribution of both pre-existing and gradually evolving *de novo* resistance. Indeed, our single cell profiling experiments revealed pre-existence of phenotypes that mapped to a major phenotypic cluster in one of the erCriz cell lines (**Fig. 1G**). Our framework is also compatible with the possibility that some mutational events lead to a dramatic fitness increase, which represents a special case within a more inclusive paradigm. On the other hand, given the evidence of the insufficiency of well-recognized “causes” of resistance such as EML-ALK amplification, single copy L1196M mutation, and alternative RTK overexpression to confer maximal resistance in the H3122 experimental model (**Fig. 4F**), it is also possible that at least in some cases, attribution of resistance to a single mechanism might be incorrect. We would like to point out that a commonly used research study algorithm, which investigates how drug sensitivity is modulated by up and down regulation of a putative resistance “driver”, followed by validation of this resistance association in clinical samples, does not discriminate between single hit and multifactorial resistance.

Finally, our data suggests that intermediate stages during the evolution of drug resistance may represent temporally restricted targets for combination or sequential therapies, using drugs that expose collateral sensitivities in evolving cells, paralleling the previously reported existence of collateral sensitivities in mutational intermediates in the evolution of resistance in Ph+ ALL^3^. Whereas the generalizability and translational potential of elevated sensitivity to lapatinib are unclear at this point, our results support the notion that adequate understanding of the forces that shape somatic evolution during the acquisition of drug resistance could open the way to designing evolutionary-informed therapy approaches, focused on staying a step ahead of resistance, rather than just reacting to it when it (inevitably) arises. Achieving progress in this direction would require dedicated multidisciplinary studies, integrating clinical studies with experimental and mathematical modeling.

## METHODS

### Derivation of Resistant Cell Lines

Parental and ALK-TKI resistant H3122 cells were grown in RPMI (Gibco, Thermo Fisher) supplemented with 10% FBS (Serum Source), penicillin/streptomycin, insulin (Gibco) and anti-clumping agent (Gibco). ALK-TKI resistant cell lines were generated by further evolving lines used in^14^, derived through a progressive increase in ALK TKI concentrations, eventually they were maintained in 0.5 µM crizotinib, 0.2 µM ceritinib, 2.0 µM alectinib and 2.0 µM lorlatinib. Alternatively, resistant lines were derived by exposing treatment naïve H3122 cells directly to the high ALK TKI concentrations, mentioned above, for 3-4 months, with re-plating every 2-4 weeks to relieve spatial growth constraints.

### Short Term Viability Assays

Short term cell viability was measured using Cell-Titer Glo reagent (Promega). Typically, 2000 cells per well were plated in a 96-well plate (Costar). Drugs were added 24h later, and the assays were performed using manufacturer recommended protocol 3-4 days after drug addition. Viability was calculated first by subtracting the luminescence of empty wells then by division by average DMSO luminescence. Dose response curves were generated by fitting the following equation to the data: 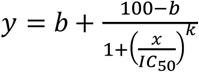; where *y* is luminescence, *x* is drug concentration, *IC*_50_ is the half maximal inhibitory concentration, *k* is the hill slope and *b* is the luminescence as the drug concentration approaches ∞.

### Clonogenic Assays

Cells were plated in the presence of ALK-TKIs or DMSO vehicle control at varying densities into 6 cm dishes or multi-well plates in duplicates or triplicates, and grown for 10 days, at which points they were fixed and stained with crystal violet, following protocols described in ^59^. To measure the evolution of gradually increasing resistance, nuclear mCherry expressing H3122 cells were plated at ∼400,000 cells per 6cm dish, allowed to attach overnight, and exposed to DMSO vehicle control, 0.5μM crizotinib or 0.5μM lorlatinib the following day. After culturing for 1-3 weeks, cells were harvested and seeded in 96-well plates (Costar) at 50 cells/well for DMSO control, or between 50 and 500 cells for crizotinib and lorlatinib. Number of colonies and colony sizes were measured 1 week later for colonies larger than 1000 pixels (∼5 cells), based on fluorescent area. To minimize the impact of variability in seeding numbers, clonogenic survival in the presence of ALK inhibitors was normalized to clonogenic data in the DMSO controls.

### Determining frequency of Resistance Initiating Cells

Cells were plated with initial seeding densities of 400, 400, 800, 1400 and 400 cells/well in DMSO (0.1%), crizotinib (0.5µM), ceritinib (0.1µM), alectinib (0.5µM) and lorlatinib (0.5µM) treated plates respectively, in a 96-well plate (Falcon). Five additional 2x dilutions were generated from these wells, each with 10 separate wells. ALK inhibitors were added after 24h. After 7 weeks of treatment, wells containing no colonies larger than ∼ 50 cells were counted. The natural log of the proportion of wells without colonies was fitted linearly against the initial cell number in each well. The number of cells per resistance - initiating cell was calculated as 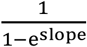 and the error of this value as 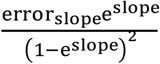.

### Clone Tracing Assay

H3122 cells were transduced at 10% efficiency (as defined by FACS analysis of dsRed expression) with ClonTracer neutral DNA barcode library^25^, kindly provided by Frank Stegmeier (Addgene #67267). Barcode containing cells were selected with puromycin and expanded for 23 days. Cells were expanded from 10^5^ cells (after puromycin selection) to 3.4×10^7^ cells. Quadruplicate cultures of 1.5×10^6^ cells each were plated into 10cm dishes in the presence of 0.1% DMSO, 0.5 µM crizotinib, 0.5 µM alectinib and 0.5 µM lorlatinib. Two pellets of 1.5×10^6^ cells were frozen to serve as an initial mixture. Cells were harvested following four weeks of cell culture. Genomic DNA was extracted using proteinase K digest, followed by phenol-chloroform purification. Barcodes were amplified, sequenced and analyzed following protocols described in ^25^ and an updated procedure provided on the Addgene website https://www.addgene.org/pooled-library/clontracer. A first round of amplifications was performed for 35 cycles using the following primers TCG ATT AGT GAA CGG ATC TCG ACG and AAG TGG ATC TCT GCT GTC CCT G for 35 cycles. Second round was performed for 15 cycles. Heatmap.2 in R (using default parameters) was used for heatmap clustering. Corrplot^60^ in R (using spearman correlation coefficients) was used to visualize correlations.

### Fluorescent *In Situ* Hybridization Analyses

Cells were cultured in 10 cm dishes in 10 ml of 1640 medium supplement with 10% FBS. After 24 hours, 100 µl of colcemid (10 µg/ml, Life Technologies, Carlsbad, CA) was added to each culture, and culture replaced to incubator for another one hour before harvesting. Cell suspension was transferred to 15 ml cortical tubes. Culture medium was removed by centrifuge at 1000 rpm for 5 min. Cell pellets were subjected to hypotonic treatment with 0.075M KCl and then fixed with Methanol and acidic acid at 3:1v/v. Cell suspensions were dropped to slides, and treated with 0.005% (wt/vol) pepsin solution for 10 min, followed by dehydration with 70%, 85%, and 100% (vol/vol) ethanol for 2 min each. Hybridization was performed by adding 10 µl of ALK dual color break apart probe on each slide (Cytocell, Cambridge UK), a coverslip was placed, and the slides were sealed with rubber cement. The specimens were subjected to denaturation at 75 °C for 3 min and hybridized at 37 °C for 16 h. The slides were washed in 0.4× saline-sodium citrate at pH 7.2 and then counterstained with DAPI. Results were analyzed on a Leica DM 5500B fluorescent microscope. Cell images were captured in both interphase and metaphase cells.

### RNAseq Analysis

Following 48 hours of culturing in the absence of inhibitors (to minimize the impact of directly induced gene expression changes), RNA was isolated for erALK-TKI and parental H3122 cells using an RNAEasy Minikit (Qiagen). Reads were generated using a MiSeq instrument. Alignment was achieved using HiSat2 and the human hg19 reference genome. Normalized reads were obtained from DeSeq2. Analysis was performed with genes with more than 25 reads in at least one sample and more than two-fold change from ALK TKI naïve cells in at least one resistant line.

### Nanostring Assay

Following 48 hours of culturing in the absence of inhibitors (to minimize the impact of directly induced gene expression changes), RNA was isolated for erALK-TKI and parental H3122 cells using an RNAEasy Minikit (Qiagen).The nCounter GX Human Cancer Reference Kit was used to determine gene reads. Genes more than one standard deviation from and more than two-fold different from the mean parental value in any sample were retained for further analysis. The 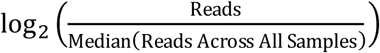 was used for all visualizations of the data. ClustVis^61^ was used to generate PCA plots. Heatmap and corresponding dendograms were generated with heatmap.2 in R, using default parameters.

### SNP Array

DNA was extracted from parental and several biological replicates of ALKi resistant H3122 cells using DNeasy Blood and Tissue Kit (Qiagen). DNA SNP sequencing was done using CytoScanHD (ThermoFisher). Reads were processed using Normal Diploid Analysis in Chromosome Analysis Suite 3.2 (ChAS, ThermoFisher).

### Single Cell Transcriptome Analysis

Single cell expression was performed using 10x Genomics platform. ∼1000 cells were analyzed per each sample, and ∼40,000 reads per cell were generated using an Illumina NextSeq 500 instrument. Demultiplexing, barcode processing, alignment, gene counting was performed using the 10X Genomics CellRanger 2.0 software. Dimension reduction was done using Uniform Manifold Approximation and Projection (UMAP)^18^, setting number of neighbors to 15. Cells were clustered using HDBSCAN on dimension reduced space^62^, with the minimum number of cells set to 30. All other parameters were set to default values.

### Gene Set Enrichment Analysis

GSEA version 4.0^63^ was used to determine the gene sets enriched in acquired resistant versus parental cell line RNA-seq profiles. We used the MSigDB Hallmark [PMID: 26771021] as the predefined gene sets and performed 10,000 permutations by gene set to determine the p-values. Gene sets with False Discovery Rate (FDR) q-value ≤ 0.25 were considered as significantly enriched.

### ChIPSeq

H3122 cells were fixed with 1% formaldehyde for 15 min and quenched with 0.125 M glycine. Chromatin was isolated by the addition of lysis buffer, followed by disruption with a Dounce homogenizer. Lysates were sonicated and the DNA sheared to an average length of 300-500 bp. Genomic DNA (Input) was prepared by treating aliquots of chromatin with RNase, proteinase K and heat for de-crosslinking, followed by ethanol precipitation. Pellets were resuspended and the resulting DNA was quantified on a NanoDrop spectrophotometer. Extrapolation to the original chromatin volume allowed quantitation of the total chromatin yield. An aliquot of chromatin (30 µg) was precleared with protein A agarose beads (Invitrogen). Genomic DNA regions of interest were isolated using 4 µg of antibody against H3K27Ac (Active Motif, cat# 39133, Lot# 01518010). Complexes were washed, eluted from the beads with SDS buffer, and subjected to RNase and proteinase K treatment. Crosslinks were reversed by incubation overnight at 65 °C, and ChIP DNA was purified by phenol-chloroform extraction and ethanol precipitation. Quantitative PCR (qPCR) reactions were carried out in triplicate on specific genomic regions using SYBR Green Supermix (Bio-Rad). The resulting signals were normalized for primer efficiency by carrying out qPCR for each primer pair using Input DNA. Illumina sequencing libraries were prepared from the ChIP and Input DNAs by the standard consecutive enzymatic steps of end-polishing, dA-addition, and adaptor ligation. After a final PCR amplification step, the resulting DNA libraries were quantified and sequenced on Illumina’s NextSeq 500 (75 nt reads, single end). Reads were aligned to the human genome (hg38) using the BWA algorithm (default settings). Duplicate reads were removed and only uniquely mapped reads (mapping quality ≥ 25) were used for further analysis. Alignments were extended in silico at their 3’-ends to a length of 200 bp, which is the average genomic fragment length in the size-selected library and assigned to 32-nt bins along the genome. <u>Comparison with</u> <u>Transcriptomics</u>. RNASeq reads were reanalyzed using HISAT2 and hg38 for proper comparison with ChIPSeq data. RseQC was used to check stranded information. HTSeq-count was used for reads counting. Raw ChIPSeq files were marked-duplicate using picard and MACS2 was used to call peaks (q<0.05). Peaks from four samples were merged using bedtools. Black list was removed. Featurecounts was used for read counting. All genes within ±20kb of a peak are used for comparisons. Peaks with less than 15 reads in all samples with ChIPSeq and RNASeq data (naïve, erCriz0, erAlec0 and erLor0) were excluded. Genes with less than 25 reads in all samples with ChIPSeq and RNASeq data were excluded. Gene and peak levels were normalized using the DESeq2 rlog function. Genes with a difference in rlog values between erTKI and naïve cells of 1 or greater are considered upregulated. Genes with a difference in rlog values between erTKI and naïve cells of -1 or less are considered downregulated. All other genes are considered neutral. Peak values that are associated with multiple genes are used multiple times. <u>Tracks</u>. MACS2 was used to call peaks. Partek was used to visualize chromosomal tracks.

### Immunoblots

Protein expression was analyzed using NuPAGE gels (ThermoFisher), following manufacturer’s protocols. The following antibodies, purchased from Cell Signaling, were used: HER2 (4290), EGFR (2963), p-ALK Y1604 (3341), ALK (3633), p-Akt S473 (4060S), Akt (9272), pERK T202/T204 (4370S), ERK (4695), p-Stat3 Y705 (9145), Stat3 (9139), p-S6 S235/S236 (4858S) and s6 (2317). Anti-β-actin antibody was purchased from Santa Cruz (47778) and used at a 1:20,000 concentration. Secondary antibodies with H+L HRP conjugates were purchased from BioRad (anti-rabbit: 170-6515, anti-mouse:170-6516). HRP chemiluminescent substrate was purchased from Millipore (WBKLS0500). Images were taken using an Amersham Imager 600 (GE Healthcare Life Sciences).

### Xenografts Studies

erALK-TKI or parental H3122 were suspended in 1:1 RPMI/Matrigel (ThermoFisher) mix and subcutaneously implanted into 4-6 week old NSG mice, with two contralateral injections per animal, containing 10^6^ tumor cells each. After 3 weeks, the animals were treated with 25mg/kg alectinib (purchased from Astatech), 6mg/kg lorlatinib (obtained from Pfizer) or vehicle control via daily oral gavage. Tumor diameters were measured weekly using electronic calipers, and tumor volumes were calculated assuming spherical shaped tumors. Tumors in the collateral sensitivity experiments were treated with 11mg/kg alectinib for one week before switching to 50mg/kg alectinib. Xenograft studies were performed in accordance with the guidelines of the IACUC of the H. Lee Moffitt Cancer Center.

### CRISPR Knockout Experiments

To produce gRNA the following DNA oligos purchased from IDT were cloned into pSpCas9(BB)-2A-Puro (PX459) V2.0 vector (Addgene Plasmid #62988), using the protocol in ^64^): CAC CGT CTC TCG GAG GAA GGA CTT G with AAA CCA AGT CCT TCC TCC GAG AGA C and CAC CGC ATC CTG CTG GAG CTC ATG G with AAA CCC ATG AGC TCC AGC AGG ATG C. H3122 cells were transfected with the above constructs using jetPrime reagent (polyPlus). 48 hours following transduction, 10^5^ cells were seeded per 6 cm dish for a clonogenic assay. To account for differences in transfection efficiency between parental and resistant cell lines, control transfections were performed using MIG GFP expressing plasmid and analyzed for percentage of GFP expressing cells using flow cytometry. Normalization was performed by multiplying the number of colonies by the transfection coefficient (ratio between fraction of GFP+ cells in the GFP control transfection between a given cell line and parental cells).

### cDNA Expression

Entry cDNA ORFs in pDONOR223 or pENTR221 were obtained from human ORFeome collection v5.1 or Life Technologies, respectively. Lentiviral expression constructs were generated by Gateway swap into pLenti6.3/V5-Dest vector (Life Technologies). Oncogenic fusion gene *EML4-ALK* variant 1 wild type or the *L1196M* point mutant was cloned into the retroviral pBabe-puro backbone^31^ (provided by J. Heuckmann, Universität zu Köln, Köln, Germany). Lentiviral and retroviral particles were produced, and used for transduction of H3122 cells following standard protocols, described in https://www.addgene.org/protocols/lentivirus-production/ and https://www.addgene.org/viral-vectors/retrovirus/retro-guide/.

## Supporting information

Mathematical supplement

## Acknowledgments

We thank M. Janiszewska, A. Goldman and A. Rozhok for their critical reading of this manuscript and discussions. I. Raplee for assistance with the RNASeq pipeline. This work was partially supported by Moffitt Lung Cancer Center of Excellence and the NIH (U54 administrative supplement 10-18279-04-13); DLP was partially funded by an IMFAHE travel fellowship. JGS is grateful to the NIH for their generous loan repayment program and to the Paul Calabresi Career Development Award for Clinical Oncology (NIH K12CA076917). This work has been supported in part by the Flow Cytometry Core, the Genomics Core and the Bioinformatics Core at the H. Lee Moffitt Cancer Center and Research Institute.

## Supplementary Figures

**Fig. S1.**
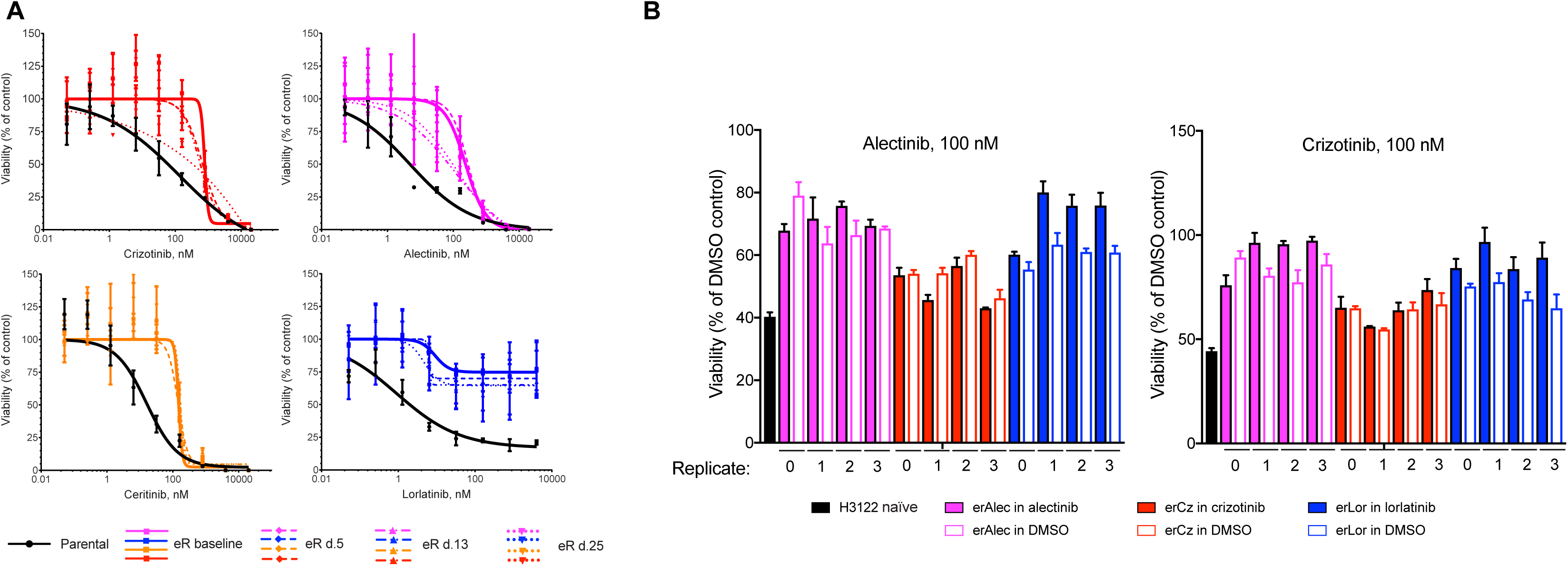
**A.** Stability of ALK-TKI resistance is assessed by analysis of ALK TKIs sensitivity of indicated erALK TKI cell lines cultured in the absence of the drugs for indicated periods of time using Cell Titer Glo assay. Error bars show standard deviation. **B.** Stability of ALK TKI resistance of independently derived cell lines, assessed by sensitivity to 100 nM alectinib and crizotinib following 4 weeks of culture in the absence of the drug. Notation of individual cell lines is identical to that used in Fig. 1F. Error bars show standard deviation.

**Fig. S2.**
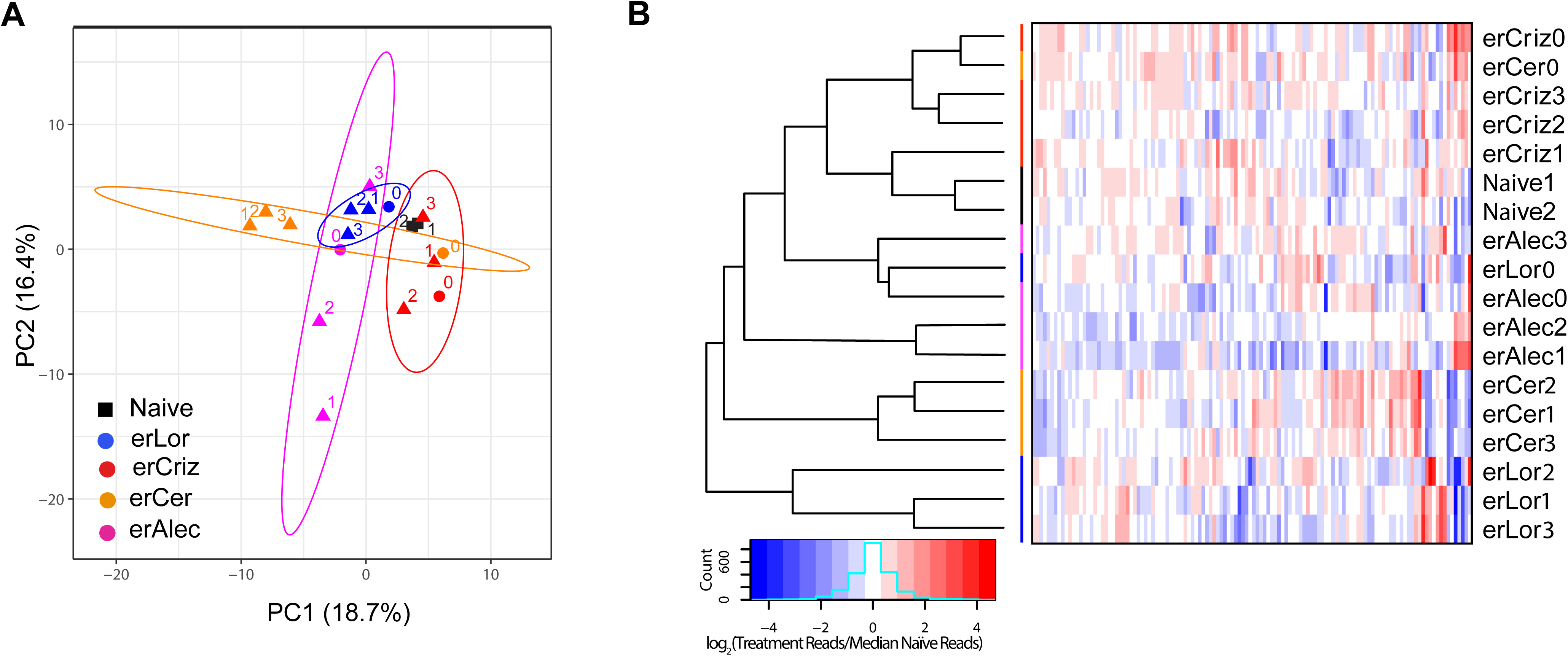
PCA analysis (**A**) and hierarchical clustering analysis (**B**) for the differentially expressed Nanostring nCounter GX human cancer reference panel transcripts of the indicated samples. Notation of individual cell lines is identical to that used in Fig. 1F.

**Fig. S3.**
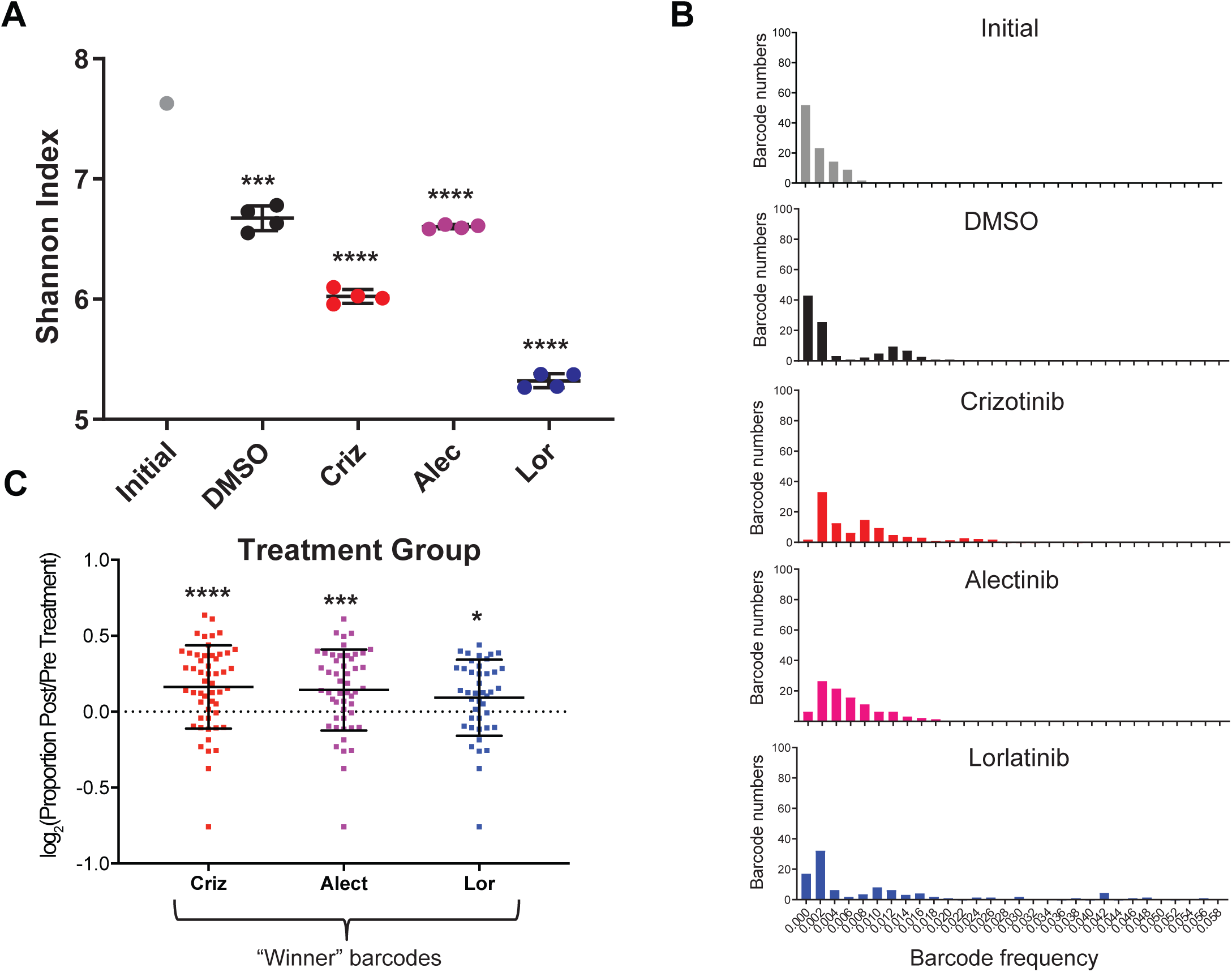
**A.** Shannon diversity of all barcodes after treatment. Error bars show standard devivation. ***, **** represent p<0.001 and p<0.0001 in a one-sample, two-tailed, t-test comparing mean Shannon diversity to the initial. Error bars show standard deviation. **B.** Distributions of clone sizes tagged with barcodes, exceeding the highest frequency pre-treatment in at least one sample. In (**A**) and (**B**) replicates in the same treatment are merged, using the mean frequency across all replicates. **C.** Frequency of barcodes under DMSO conditions. Only barcodes exceeding the highest frequency in the initial mixture are shown. Error bars show standard deviation. *, *** and **** represent p<0.05, p<0.001 and p<0.0001 in a one sample, two-tailed, t-test comparing means to zero.

**Fig. S4.**
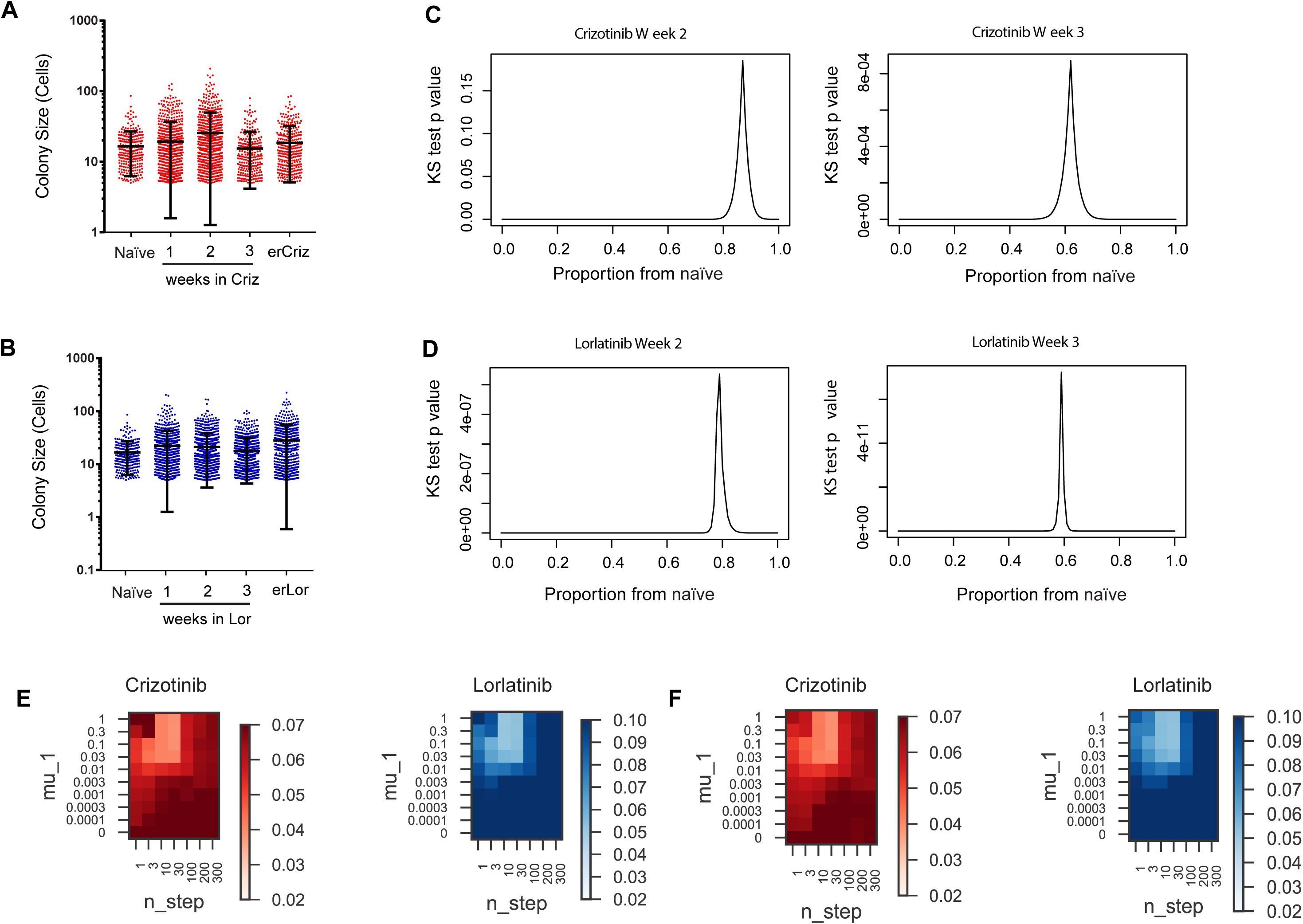
**A**, **B**. DMSO control colony size distribution of treatment naïve H3122 cells, cultured in the presence of 0.5 µM crizotinib (**A**) or lorlatinib (**B**) for the indicated amount of time prior to the clonogenic assay. Error bars show standard deviation. **C-D**. Kolmogorov-Smirnov test estimate for likelihood that the distribution at an intermediate time point can be explained by combined sampling from DT cells (naïve cell colony distribution) and erALK-TKI cells, for growth in crizotinib (**C**) and in lorlatinib (**D**). **E, F** Kullback–Leibler divergence-based comparison of the experimental data with the outcomes of simulations, covering parameter spaces for the indicated mutation probabilities and number of mutational steps, with the inclusion of death probability (**E**) and an additional inclusion of mutations that cause bi-directional fitness changes (**F**).

**Fig. S5.**
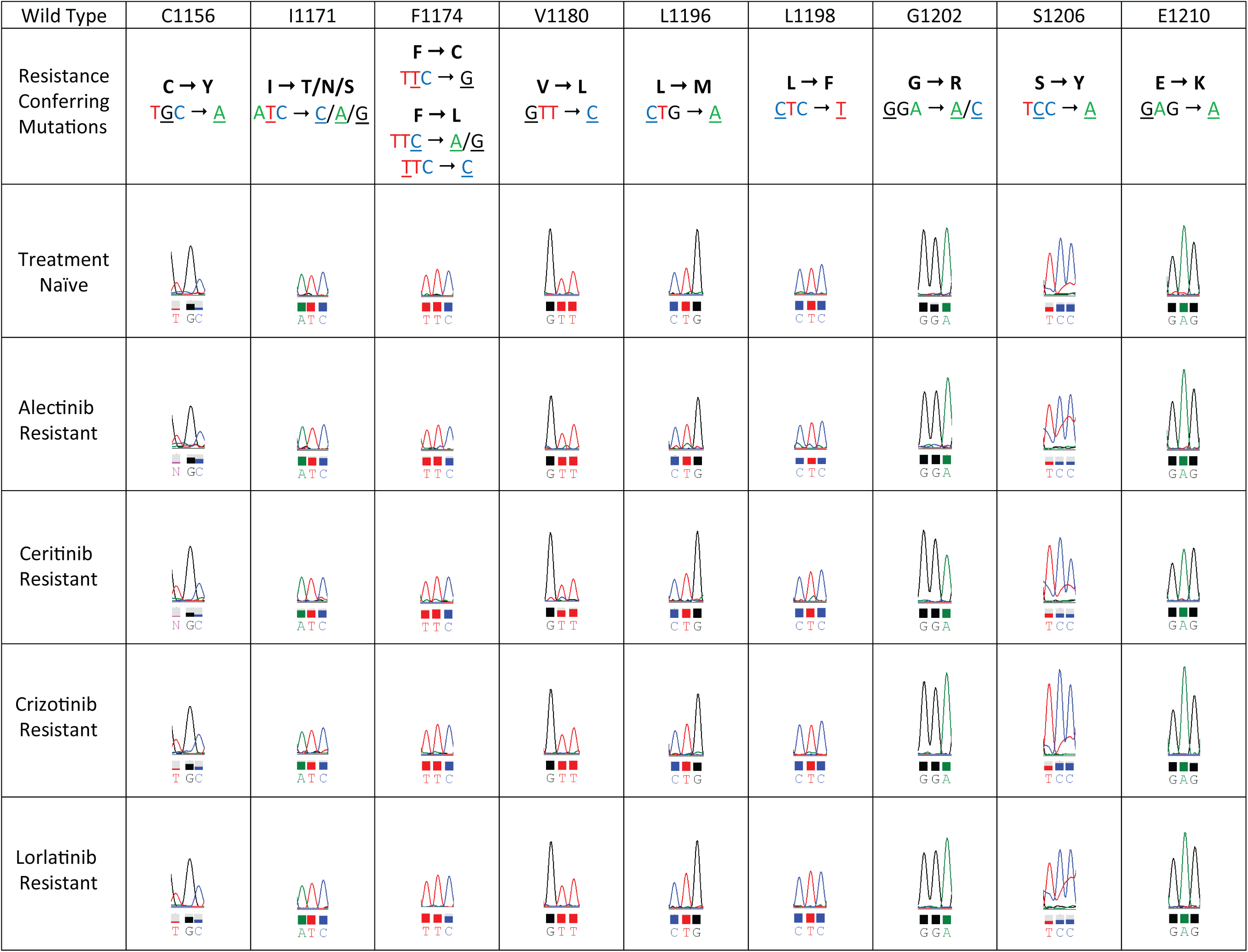
Sanger sequencing traces for the indicated resistance-associated EML4-ALK hotspot mutations.

**Fig. S6.**
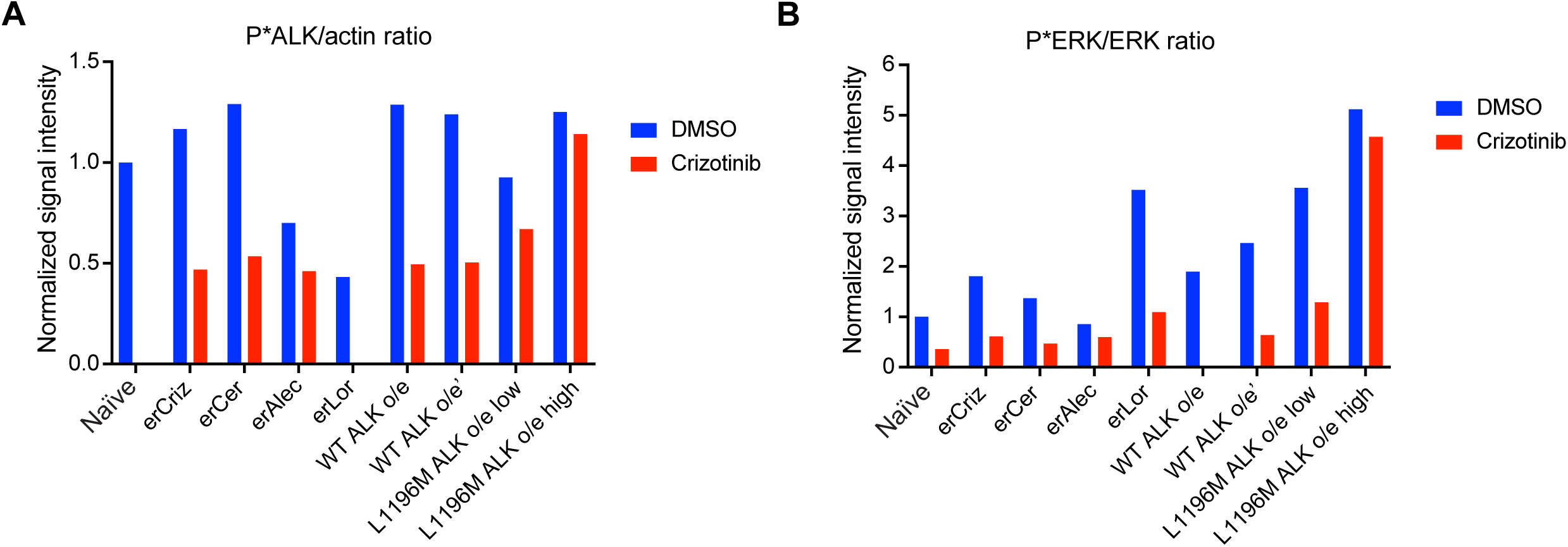
Quantitation of immunoblot data from Fig. 4E. **A**. Phospho-EML4-ALK/actin ratio. To account for changes in EML4-ALK expression in overexpressing cell lines, the data is normalized to EML4-ALK/actin ratio in the DMSO control naïve cells. **B**. Phosphorylated to total ERK ratio, normalized to the ratio in the DMSO control naïve cells.

**Fig. S7.**
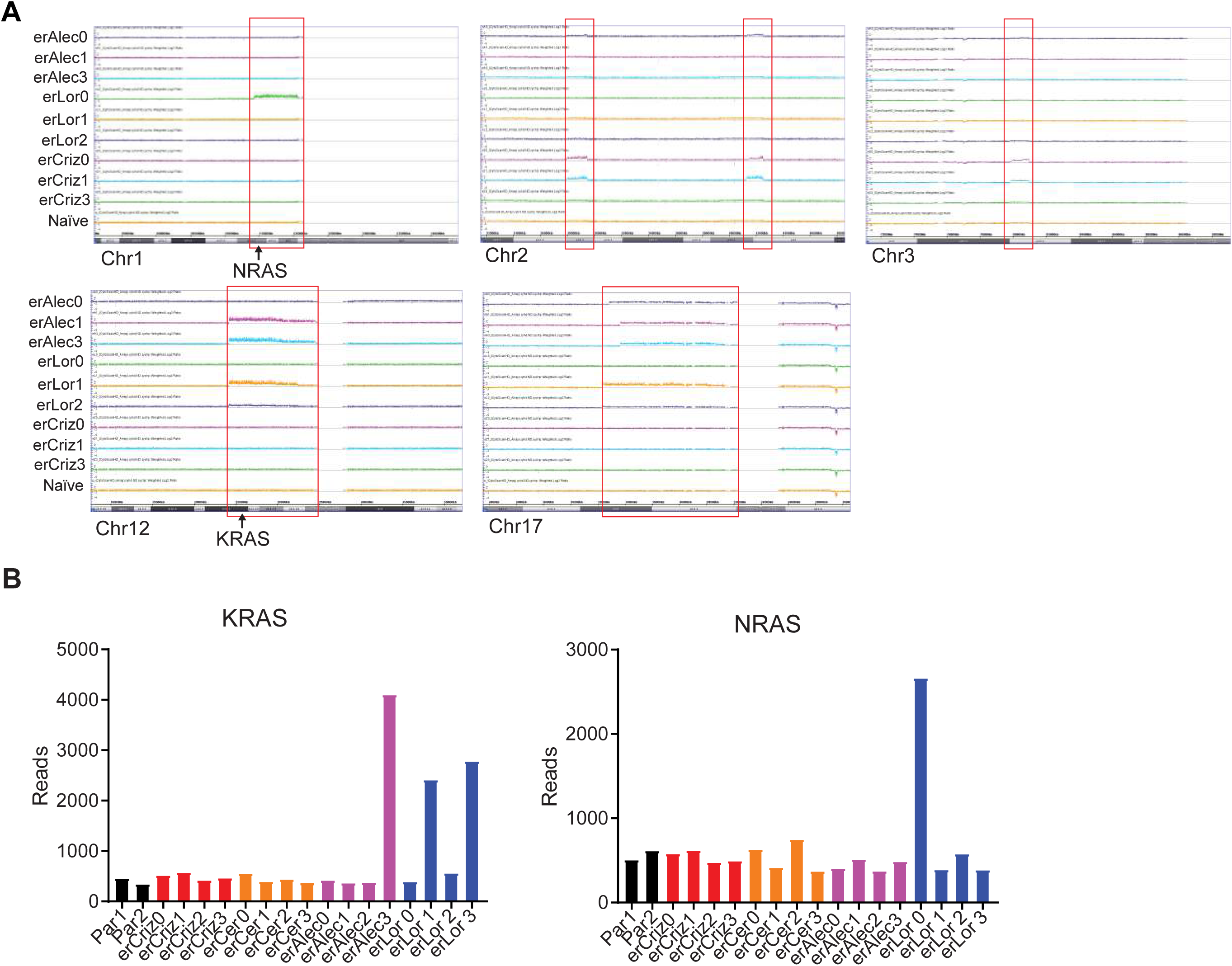
**A.** Weighted log_2_ ratio in regions selected based on differences between parental H3122 and at least one resistant line, differences are highlighted by red boxes. The genomic locations of KRAS and NRAS are highlighted. **B.** KRAS and NRAS reads obtained from NanoString expression data.

**Fig. S8.**
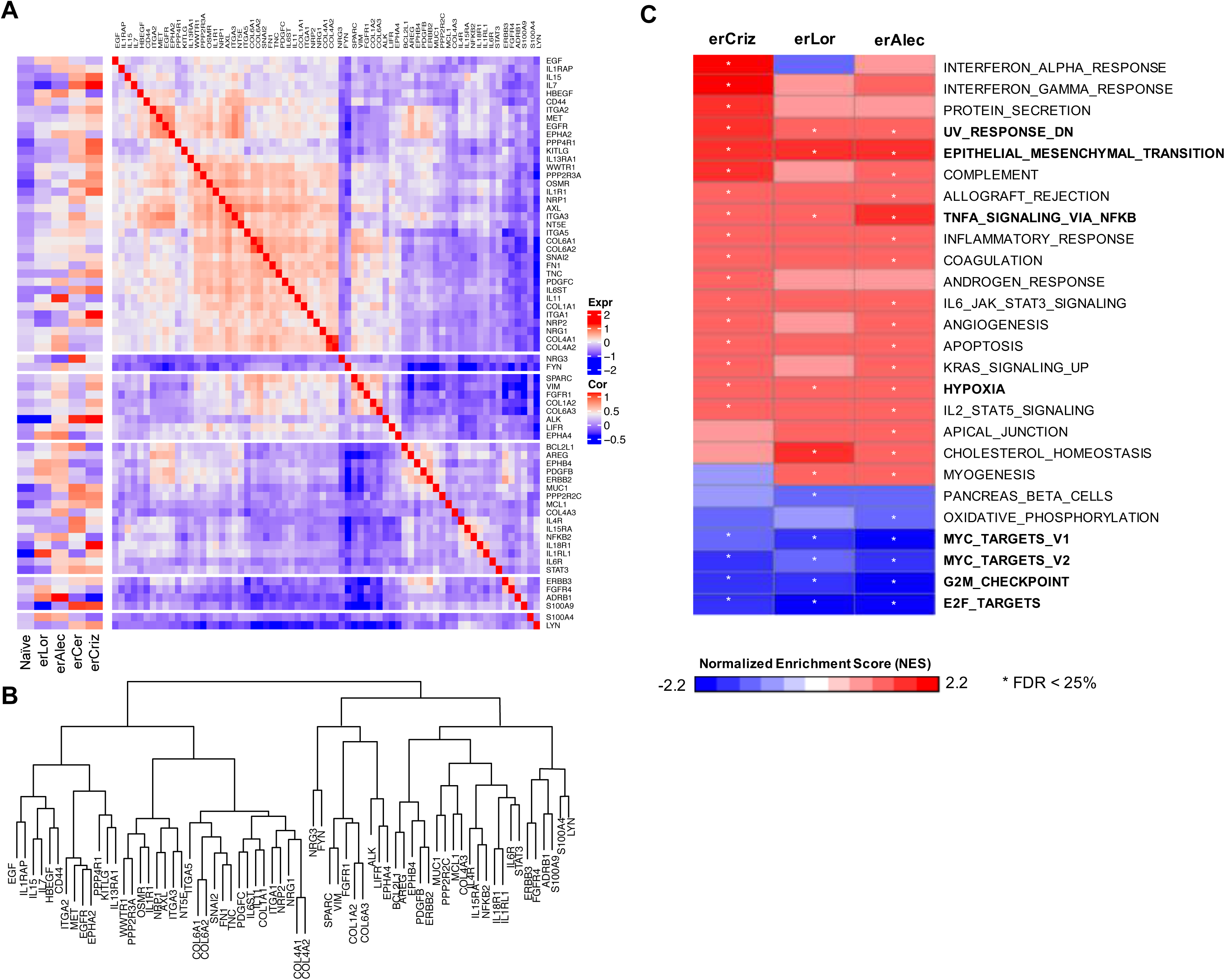
**A**. Correlation analysis for co-expression of resistance-associated genes, upregulated in erALK-TKI cell lines. **B**. Hierarchical clustering dendrogram for the individual overexpressed genes. **C.** GSEA results from three acquired resistant cell lines were plotted as a heatmap using Normalized Enrichment Score (NES), where gene sets with FDR ≤ 0.25 is indicated as an asterisk (*).

**Fig. S9.**
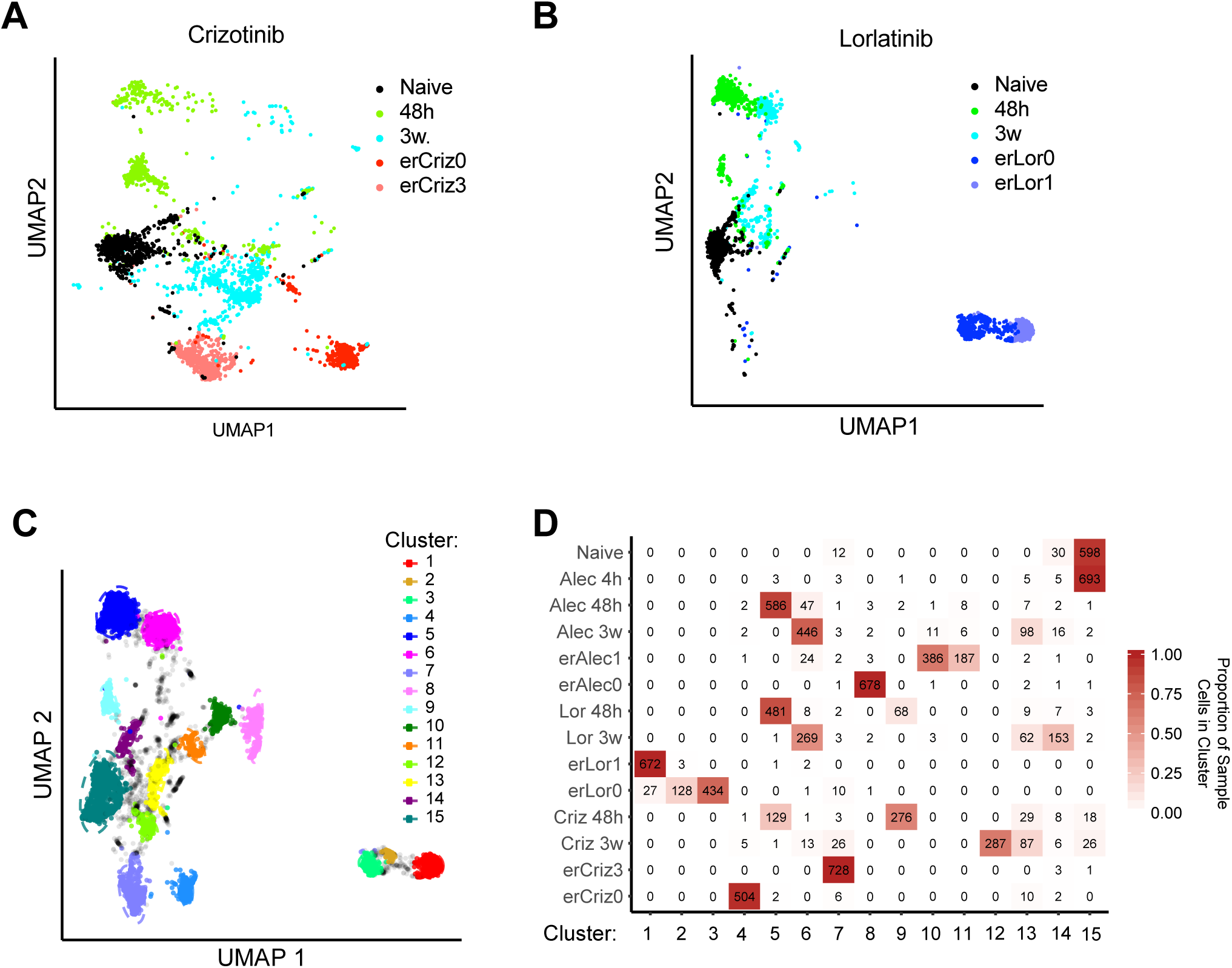
**A, B.** Single cell RNA seq based UMAP analysis of phenotypes at indicated time points post initiation of crizotinib (**A**) and lorlatinib (**B**) treatment, and in the indicated erALK TKI cell lines (one outlier cell lies outside the axes). **C.** Cluster assignment for the UMAP of single cell transcriptomics data from all samples shown in **Fig. 1G, 5B, S9 A, B**. Gray cells are unassigned. Ellipsoids show 95% confidence intervals. **D.** Heatmap reporting the number of cells in each cluster and sample. Shading is based on the proportion of cells in a sample that are in a specific cluster. Cluster-0 are unassigned cells.

**Fig. S10.**
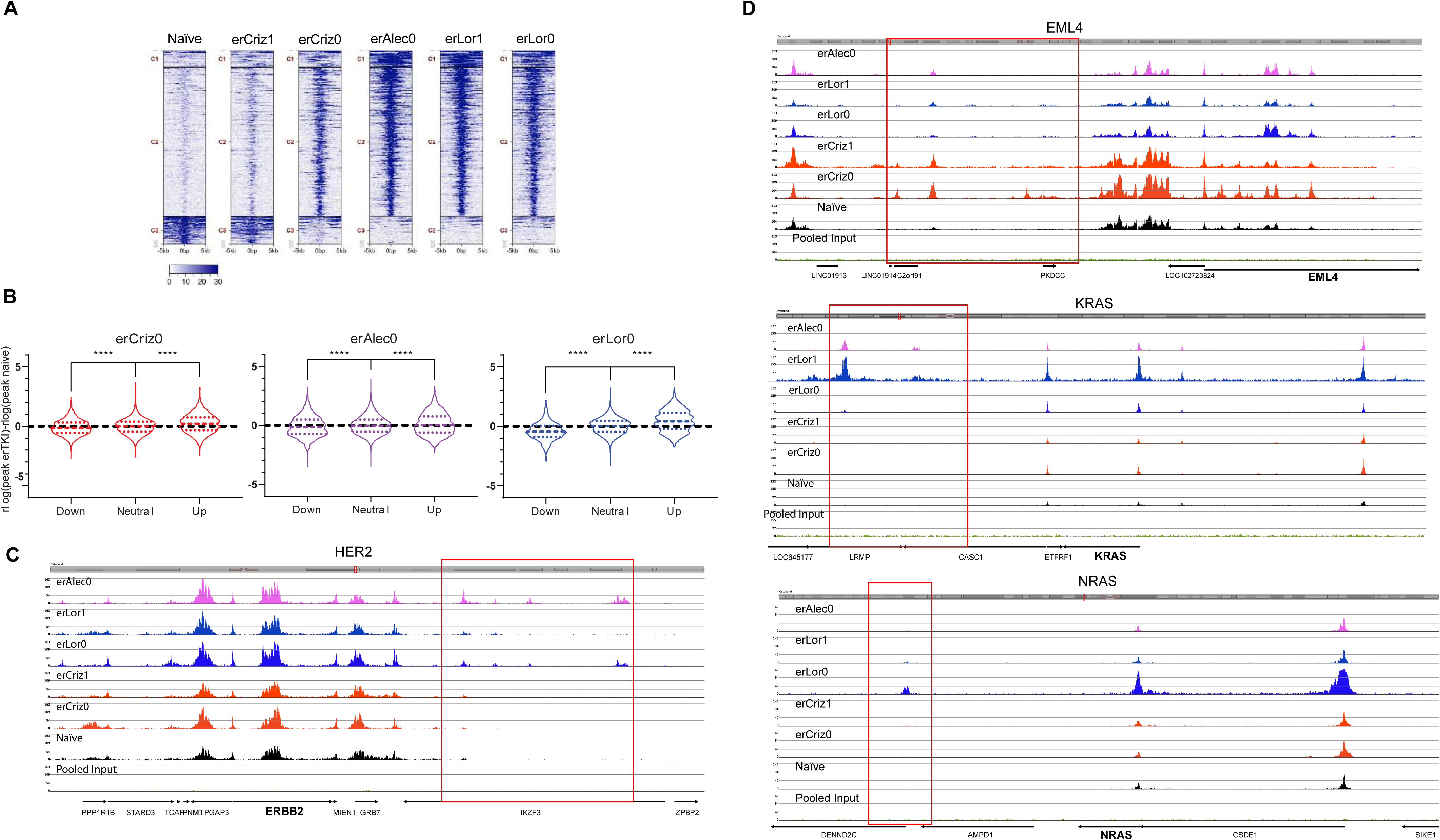
**A.** Clustered H3K27 acetylation of differentially acetylated peaks in the indicated cell lines. **B.** Changes in H3K27 acetylation in differentially expressed genes. Black dashed line represents zero, colored dashed lines are medians, colored dotted lines represent quartiles. **** represents p<0.0001 in a Kruskal-Wallis test. **C**. Elevated expression of HER2 is associated with formation of novel H3K27 peaks in the gene’s vicinity. **D**. Changes in H3K27 acetylation in the vicinity of the indicated genes.

**Fig. S11.**
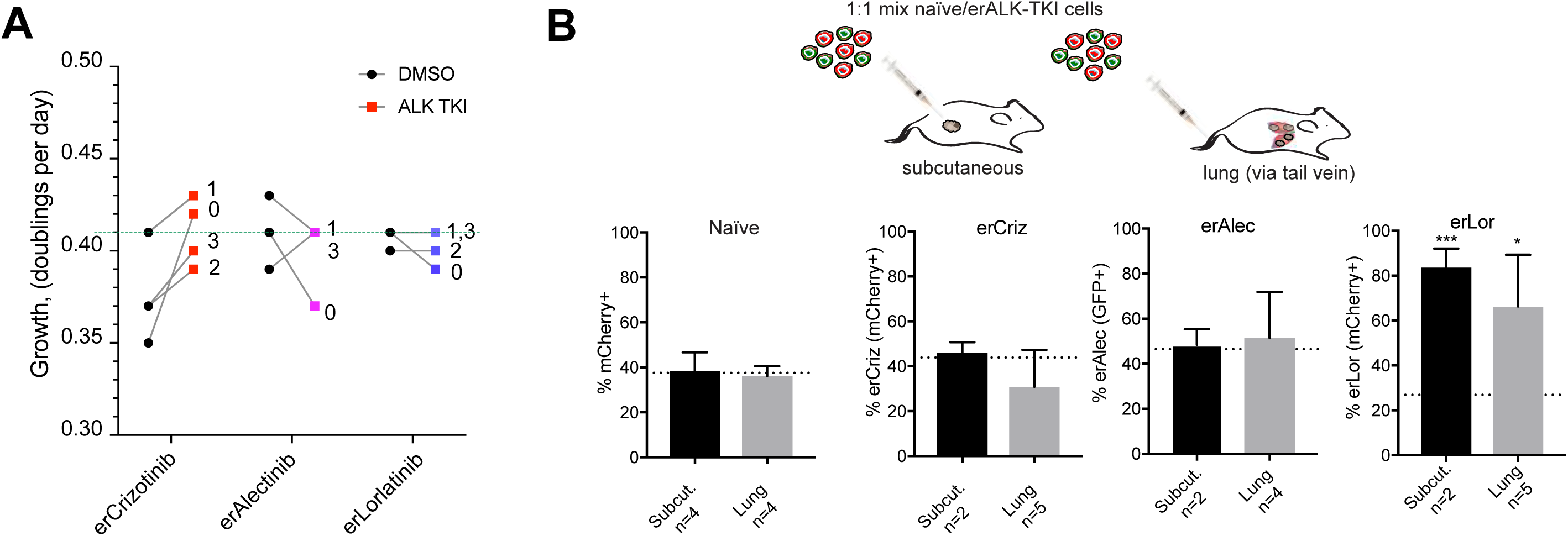
**A.** Net growth rates of the indicated erALK-TKI cell line in the presence and absence of 0.5 µM of the indicated ALK inhibitors, determined by counting cells over 4 weekly passages. Dashed line represents growth rates of therapy naïve cells cultured under DMSO control. **B**. Flow cytometry-based quantitation of the frequency of the indicated erALK TKI cells (evolved through dose escalation), transplanted while mixed with differentially labelled therapy naïve H3122 cells subcutaneously or orthotopically, via tail vein injection. Dashed lines represent frequency of cells in the initial mixtures. * and *** represent p<0.05 and p<0.001 respectively in a one-sample, two-tailed, t-test comparing means to the initial proportion of cells. Error bars show standard deviation.

**Fig. S12.**
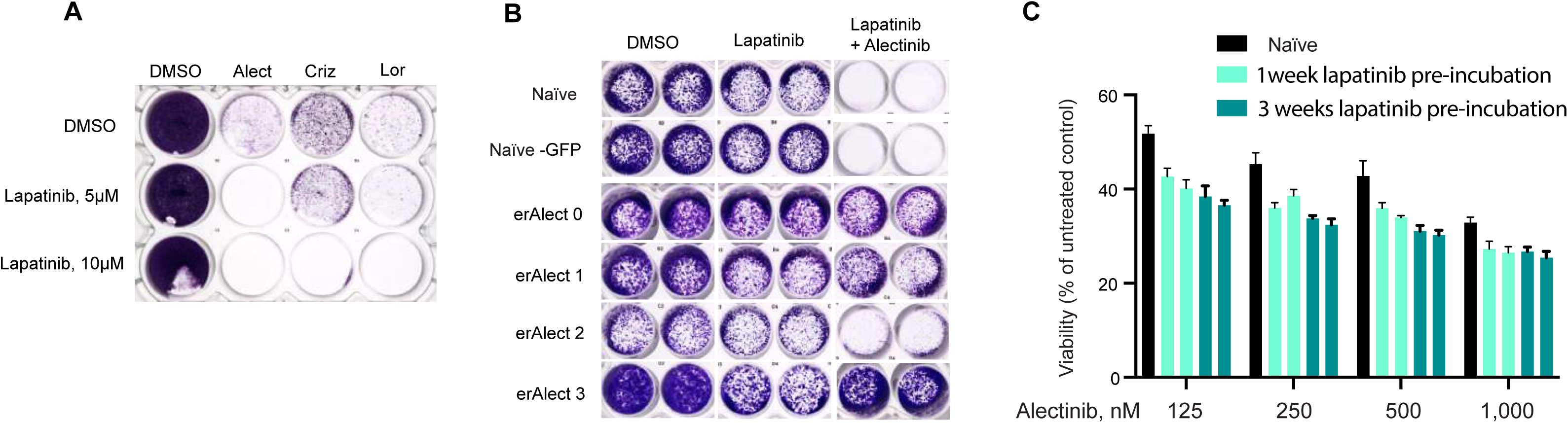
**A**. Drug naïve H3122 cells were cultured in the presence of the indicated ALK TKI (0.5 µM) or DMSO control with or without indicated concentrations of lapatinib for 6 weeks, then stained with crystal violet. **B**. Sensitivity of treatment naïve and er-ALK TKI to lapatinib mono or combined therapy, determined after 7 days of treatment by staining with crystal violet. **C.** Alectinib sensitivity of H3122 cells, cultured in the presence of 10 µM lapatinib for 1 and 3 weeks, was compared to the sensitivity of naïve H3122 cells using Cell Titer Glo viability assay. Error bars show standard deviation.

