## Supplementary material for "Resistance to targeted therapies as a multifactorial, gradual adaptation to inhibitor specific selective pressures": Mathematical supplement

### Appendix: simulation model

#### Modeling setup

We have developed a stochastic simulation to study, *in silico*, the *in vitro* cell culture experiment (**Fig. 3A**) discussed in Results Section III of the main text. We have chosen a cellular automaton framework, an individual based model in which each individual cell is tracked through time and space, and which follows simple update rules after preset time steps ($\Delta t = 16hrs$, therefore $1\ wk \approx 9\Delta t$). Each update includes randomly chosen cellular events, which include divisions, mutations and interactions with other cells in a regular grid in 2-dimensional space (illustrated in **Math Supp. Fig. 1**).

We chose to model each colony individually, and therefore our cellular automaton model starts with a population size of one, and in each time step, each cell is assumed to have a chance to divide if there is an empty space in its (Moore) neighborhood. If it divides, it places a daughter into its current position and a uniformly randomly chosen empty neighboring space. If there is no direct empty neighboring space, it can still divide if there is an empty spot one lattice step beyond its immediate neighborhood by 'pushing' an adjacent cell to occupy that spot to divide. However, we assumed cells cannot divide by pushing two or more layers.

There are 4 parameters involved, (i-ii) the upper and lower limits of division rates, $\boldsymbol{d_0}$ and $\boldsymbol{d_1}$, (iii) the number of mutation steps taken to reach $d_1$ when starting from $d_0$, $\boldsymbol{n_{step}}$ (illustrated in **Math Supp. Fig. 2**), and (iv) mutation rate, $\boldsymbol{\mu}$. The two biological hypotheses were tested using different $n_{steps}$: $n_{step} = 1$ for a single mutation step, and $n_{step} > 1$ for multiple steps. Then, given a specific range of division rates, $[d_0, d_1]$, the mutational increase in division rate (fitness) is $\Delta d = (d_1 - d_0)/n_{step}$ (see **Math Supp. Fig. 1** for a visual explanations of $\Delta d$ in terms of other parameters).

Additionally, in our study, higher values of $\mu$ have been paired with scenarios with higher $n_{step}$ values, as such scenarios require more mutations to achieve equivalent (observed) division rates, necessitating higher mutation probabilities. The rule we used to choose $\mu$ for $n_{step}$ is $\mu_{n_{step}} = n_{step} * \mu_1$, where $\mu_1$ is the mutation rate in the single step scenario. In this way, we model low probability (rare) mutations with huge fitness effects to high probability (common) mutations with tiny fitness effects.

The biological stochasticity is accounted for by randomly deciding the (i) spot to divide, as well as the (ii-iii) timing of division and mutation events, occurring with probability $d$ and $\mu$ respectively. **Math Supp. Fig. 1** shows a detailed algorithm for this model. With the inputs being division rate ($d^{(i)}$) and mutation stage ($0 \leq n^{(i)} \leq n_{step}$) of a cell at the $i$-th time step, if the cell divides, updated $d^{(i+1)}$ and $n^{(i+1)}$ are outputs, as well as a new topology of cell spatial occupation.

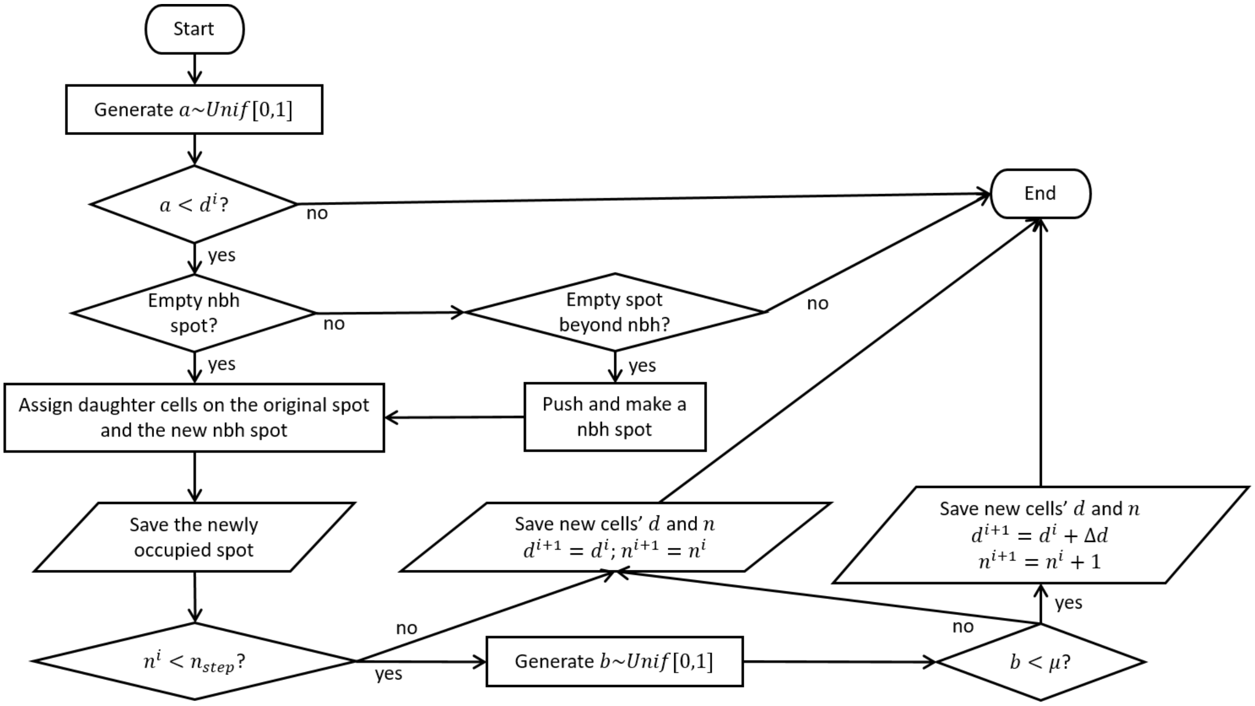


*Figure 1. Algorithm to update each cell into the next time step in the cellular automaton model.*

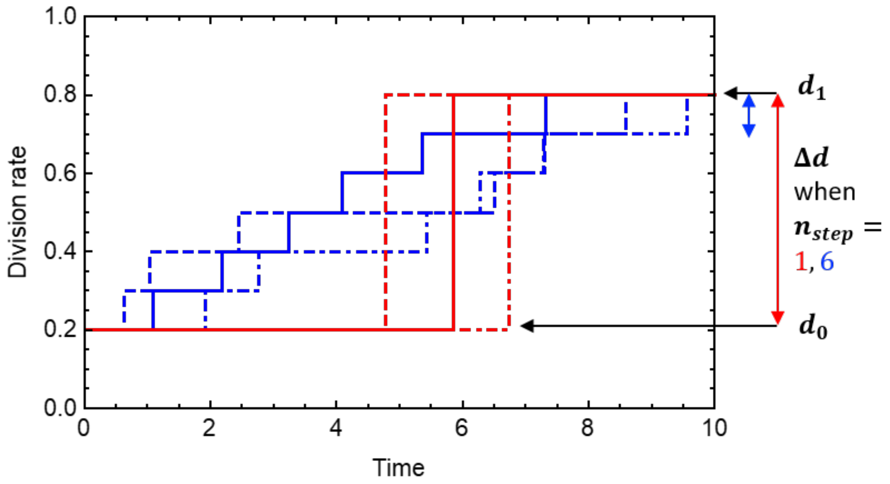


*Figure 2. Illustration of the evolutionary increase in division rate, randomly increasing (by mutation) over time. With a given range of division rate, $[d_0, d_1]$, three examples of fitness histories are shown for each scenario: the single-step scenario (red curves), and the six-mutation step scenario (blue curves).*

#### Parameter calibration

We calibrated model parameters in two steps.

**(Step 1) calibration of range of division rates $[d_0, d_1]$:**

To calibrate the initial division rate, $d_0$, we utilized the distribution data from the control assay colony sizes (**Fig. 3C**). Assuming only minor variance in the distribution of drug naïve cell division rates (in the presence of drug), we calibrated a single fixed value of $d_0$ using the best fit to the observed initial distribution. However, when calibrating the upper limit, we derived multiple values of $d_1$s, each of which is relevant to an individual resistant colony size. This empirical distribution of varying $d_1$s was used to generate diverse *in silico* colonies. Specifically, we first generated 500 *in-silico* colonies for every $d_1 \in \{0.01, 0.02, 0.03, \ldots ,0.99,1.0\}$ (with mutation not allowed for fully resistant cells, $d = d_1$), based on the algorithm described in **Math Supp. Fig. 1**. Then we interpolated values of $d_1$ and the median of 500 colony sizes in $d_1$, $\{d_1, \widetilde{P_{d_1}}\}$. These values were used to parameterize $d_1$ (**Math Supp. 3A**) for each colony size. **Math Supp 3B** shows the distribution of $d_1$ derived in this way.

(a) (b)

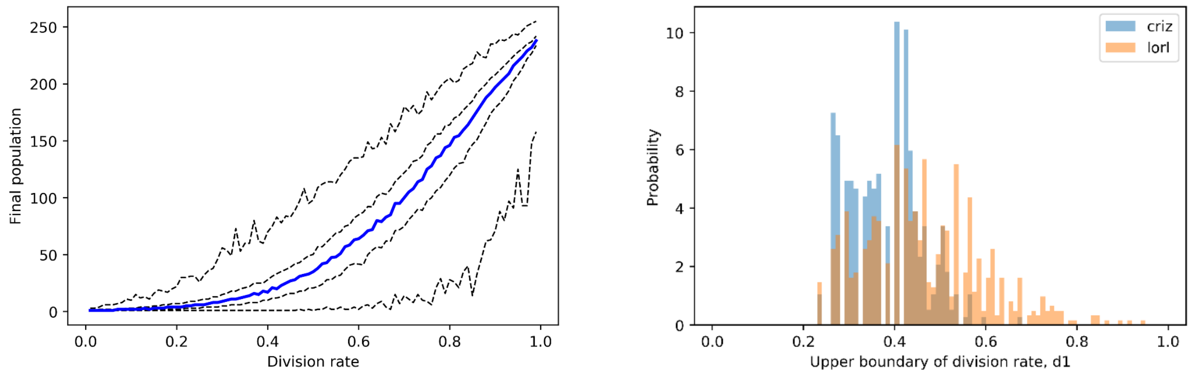


*Figure 3. (a) Simulated relationship between fixed division rates and final population sizes, after growing for one week. The thick blue line represents the median and dashed lines represent 0%, 25%, 50%, 75% and 100% quantiles of500 realizations. (b) Distributions of the upper limit of division rates, $d_1$, calibrated based on the median curve from (a).*

Based on the relative homogeneity of drug naïve cell colony sizes, in drug, we calibrated just one fixed value for $\mathrm{d}_0$. The colony sizes simulated with a fixed $\mathrm{d}_0$ show a good fit to our data (see the first couple of columns in **Math Supp. Fig. 4**). However, the colony sizes generated with a distribution of varying $\mathrm{d}_0$ (based on the median curve calibration of **Math Supp. 3A**) does not, probably due to the right skewness of the distribution for low division rates (see the second couple of columns in **Math Supp Fig 4**). Therefore, the calibration method using various division rates seems proper only for evaluating $\mathrm{d}_1$ (see the third couple of columns in **Math Supp. Fig. 4**).

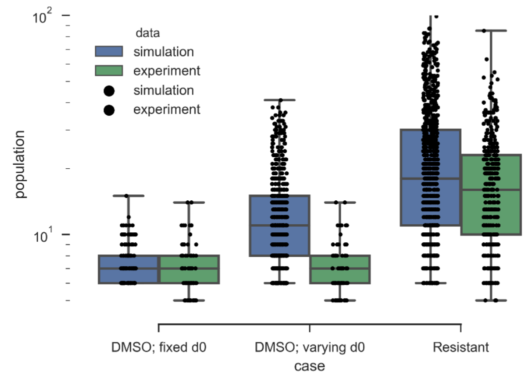


*Figure 4. Comparison between experimental data and in-silico data simulated with the calibrated parameters, $d_0$ and $d_1$.*

**(Step 2) calibration of mutation steps ($n_{step}$) and mutation rates ($\mu$)**:

Based on the calibrated $d_0$, $d_1$ and the clonogenic assay data for cells with different exposure times to the drugs (i.e., 1-, 2-, 3-weeks exposure duration), we performed an exhaustive parameter sensitivity analysis for $n_{step}$ and $\mu$ values.

For every choice of $\{n_{step}, \mu\}$, we ran 100,000 random simulations, following our stochastic algorithm, designed to mimic our experimental protocol. The simulation results in 100,000 virtual colonies of varying size when seeding cells with 1-, 2- and 3-weeks of drug exposure, as in the experimental conditions. To measure the closeness between our experimental and simulated results, we defined an error estimator between two distributions based on the Kullback–Leibler (KL) divergence. Specifically, we binned empirical observations into 10 bins based on the minimum ($m$) and maximum ($M$) values of the dataset. The lengths of the bins are all same: $\Delta b = (M - m)/9$, and the bins are half-open intervals $[m + (i-1)\Delta b, m + i\,\Delta b)$ for $i = 1, 2, \ldots, 10$. With such bin ends, we evaluated normalized bin counts for empirical data ($p_i$, $\sum_{i=1}^{10} p_i = 1$). Similarly, we binned the corresponding simulated data ($q_i$), but clipped it into the interval $[m, M]$ before binning so that the total probability equals 1 ($\sum_{i=1}^{10} q_i = 1$) (see **Math Supp. Fig. 5** for a visual explanation based on an example). As we have the data for three different exposure periods ($wk = 1,2,3$), we measured KL divergence for each week, and used the average of them as the error, which is expressed by:

$$KL = \frac{1}{3}\sum_{wk=1}^{3} \sum_{i=1}^{10} p_i^{wk}\, log\left(\frac{p_i^{wk}}{q_i^{wk}}\right).$$

We estimated the error over a range of $\{n_{step}, \mu\}$ parameters. See the main text for results.

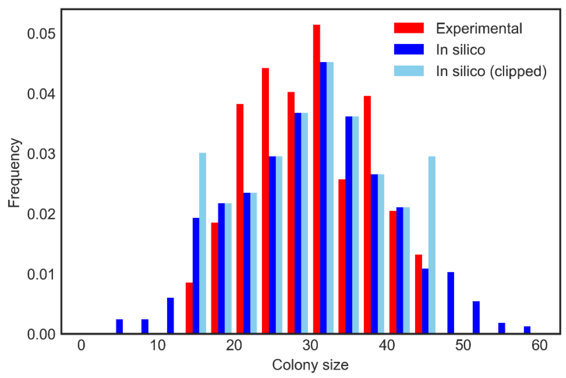


*Figure 5. An example distribution of experimental data binned into 10 bins (red), and the corresponding simulated distribution (blue) clipped by the minimum and maximum of the bin ranges (sky blue).*

#### Model extension by including (i) stochasticity in mutation step ($\Delta d$) and (ii) cell death

In the model described above, we accounted only for components that are indispensable when answering our primary question. In the main body of our article, we showed results based on this basic model. However, as an extension of this work, we developed extended models to account for two additional biological features, (i) variation in the influence of a mutation on fitness and/or (ii) cell death.

In one of the models, we assigned a death probability of $\delta = 0.01$ ($\approx 0.1\ d_0$). Therefore, each cell can die with a probability $\delta$, divide with $d$, and remain the same otherwise (M1). In the second model, in addition to a death probability ($\delta = 0.01$), we considered bi-directional random changes in proliferation rates as a result of mutations (M2). Instead of the deterministic increase in division rate $\Delta d$, we used stochastic steps $U(\{-1,1\}) \times Pois(1) \times \Delta d$, where $U(\{-1,1\})$ which follow a discrete uniform distribution drawing -1 and 1 with equal chances and arise according to a Poisson distribution with a mean of 1 timestep.

With the modified versions of the model, we carried out an identical parameter calibration and KL divergence evaluation as with earlier models. The results still support a multi-step model of resistance evolution (**Fig. S4E,F**)
